# Disrupted callosal connectivity underlies long-lasting sensory-motor deficits in an NMDA receptor antibody encephalitis mouse model

**DOI:** 10.1101/2022.09.29.510196

**Authors:** Jing Zhou, Ariele L. Greenfield, Rita Loudermilk, Christopher M. Bartley, Chun Chen, Xiumin Chen, Morgane Leroux, Yujun Lu, Deanna Necula, Thomas T. Ngo, Baouyen T. Tran, Patrick Honma, Kelli Lauderdale, Chao Zhao, Xiaoyuan Zhou, Hong Wang, Roger A. Nicoll, Cong Wang, Jeanne T. Paz, Jorge J. Palop, Michael R. Wilson, Samuel J. Pleasure

**Affiliations:** Department of Neurology, University of California, San Francisco, San Francisco, CA 94143, USA; Weill Institute for Neurosciences, University of California, San Francisco, San Francisco, CA 94143, USA; Center for Encephalitis and Meningitis, University of California, San Francisco, San Francisco, CA 94143, USA; Department of Psychiatry and Behavioral Sciences, University of California, San Francisco, San Francisco, CA 94143, USA; Gladstone Institute of Neurological Disease, San Francisco, CA 94158, USA; Department of Neurology and Institute of Neuroscience of Soochow University, Second Affiliated Hospital of Soochow University, Suzhou, 215004, China; Department of Cellular and Molecular Pharmacology, University of California, San Francisco, CA 94158, USA; School of Acupuncture Moxibustion and Tuina, Shanghai University of Traditional Chinese Medicine Intelligent Rehabilitation, Pudong New Area, Shanghai, 201203, China; Neuroscience Graduate Program, University of California, San Francisco, San Francisco, CA 94158, USA; Center for Data Driven Discovery in Biomedicine, Children’s Hospital of Philadelphia, Philadelphia, Pennsylvania, PA 19146, USA; Department of Physiology, University of California, San Francisco, CA 94158; Division of Membrane Physiology, Department of Molecular and Cellular Physiology; Department of Rehabilitation Medicine, Tong Ren Hospital, Shanghai Jiao Tong University School of Medicine, Shanghai, 200336, China; Yuanshen Rehabilitation Institute, Shanghai Jiao Tong University School of Medicine, Shanghai, 200025, China; Queensland Brain Institute, The University of Queensland, St Lucia, Brisbane, Australia, 4072; Programs in Neuroscience and Developmental Stem Cell Biology, Eli and Edythe Broad Center of Regeneration Medicine and Stem Cell Research, Kavli Institute for Fundamental Neuroscience, San Francisco, CA 94143, USA

**Author notes:** Address correspondence to: Samuel J. Pleasure, 675 Nelson Rising Lane, #214, San Francisco, CA 94158, email address; Jing Zhou, 675 Nelson Rising Lane, NS260, San Francisco, CA 94158. equal contributions (Ariele L. Greenfield, Rita Loudermilk, Christopher M. Bartley).

## Abstract

NMDA receptor mediated autoimmune encephalitis (NMDAR-AE) frequently results in persistent sensory-motor deficits, especially in children, yet the underlying mechanisms remain unclear. This study investigated the long-term effects of exposure to a patient-derived GluN1-specific monoclonal antibody (mAb) during a critical developmental period (from postnatal day 3 to day 12) in mice. We observed long-lasting sensory-motor deficits characteristic of NMDAR-AE, along with permanent changes in callosal axons within the primary somatosensory cortex (S1) in adulthood, including increased terminal branch complexity. This complexity was associated with paroxysmal recruitment of neurons in S1 in response to callosal stimulation. Particularly during complex motor tasks, mAb3-treated mice exhibited significantly reduced inter-hemispheric functional connectivity between S1 regions, consistent with pronounced sensory-motor behavioral deficits. These findings suggest that transient exposure to anti-GluN1 mAb during a critical developmental window may lead to irreversible morphological and functional changes in callosal axons, which could significantly impair sensory-motor integration and contribute to long-lasting sensory-motor deficits. Our study establishes a new model of NMDAR-AE and identifies novel cellular and network-level mechanisms underlying persistent sensory-motor deficits in this context. These insights lay the foundation for future research into molecular mechanisms and the development of targeted therapeutic interventions.

**Graphical Abstract:** 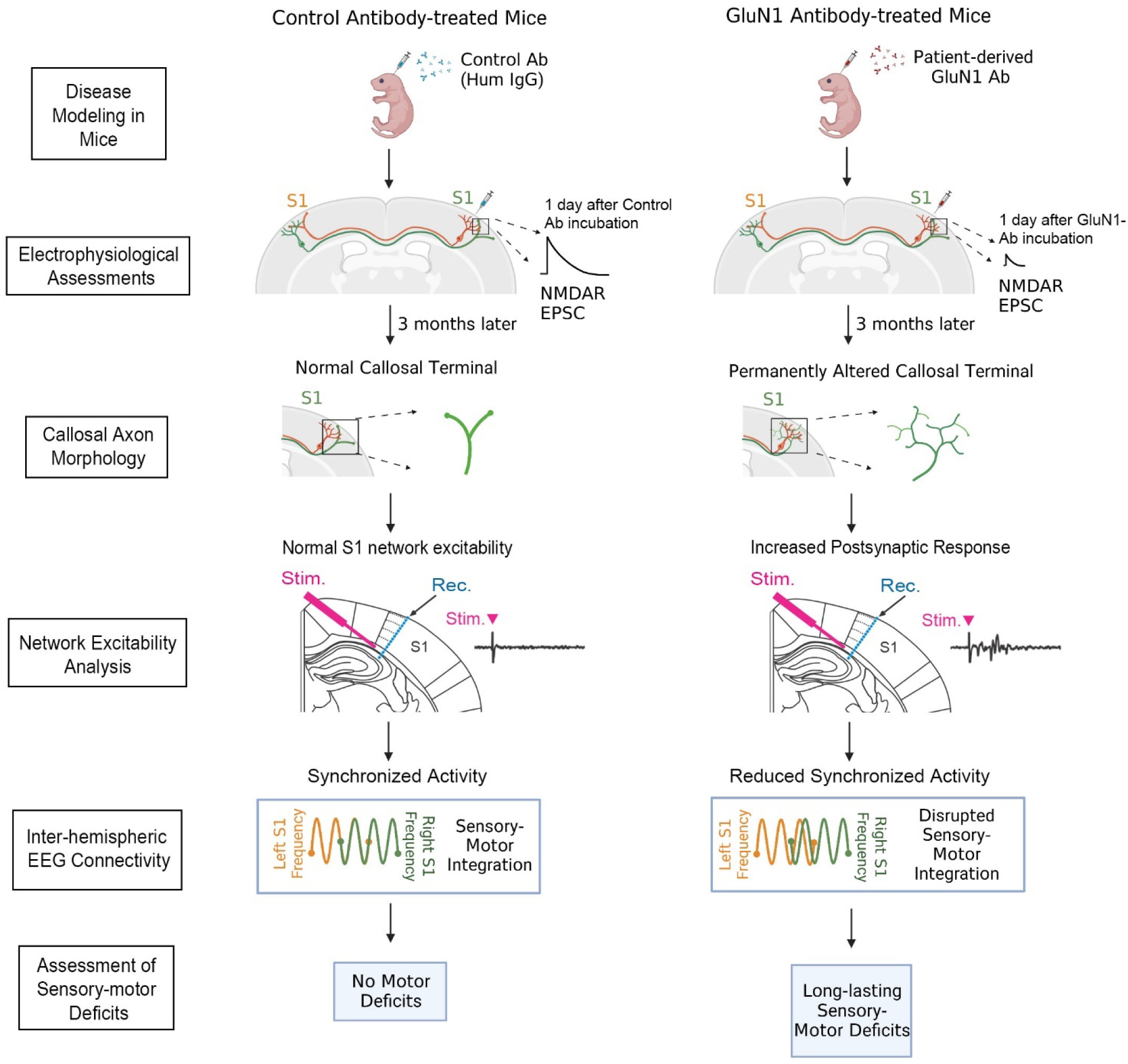

## Introduction

N-methyl-D-aspartate (NMDA) receptor mediated autoimmune encephalitis (NMDAR-AE), the most common form of autoimmune encephalitis affects both children and young adults (1, 2). Typically, NMDAR-AE is associated with antibodies against the extracellular domain of the GluN1 subunit of the NMDA receptor. Patients exhibit a range of symptoms including abnormal behaviors, psychotic symptoms, memory impairment, movement disorders, and seizures, with the severity of these symptoms varying with age (3–6).

Existing mouse models of NMDAR-AE have primarily focused on the acute behavioral deficits caused by anti-NMDAR antibodies (7–10). These autoantibody models led to a selective and reversible decrease in NMDAR surface density and synaptic localization (11–13). Accordingly, the behavioral deficits observed in these models, such as memory impairment, are reversible following the removal of the antibodies and restoration of synaptic NMDAR levels (7).

While many symptoms are responsive to immunotherapy, persistent behavioral deficits remain in many patients (14–16). Such persistent behavioral deficits are not only seen in NMDAR-AE but are also common sequela in many other autoimmune neurological disorders (17, 18). Clinical evidence in pediatric NMDAR-AE patients suggests that early onset of disease and treatment delays exceeding 4 weeks are correlated with a more severe disease trajectory, characterized by an increased risk of long-lasting sensory-motor deficits (19). Animal studies further corroborate these clinical observations, showing that transplacental transfer of anti-NMDAR antibodies can cause behavioral changes in offspring, although the underlying mechanisms of these changes are unknown (20–22). This confluence of clinical and animal research suggests that early exposure to anti-NMDAR autoantibodies may disrupt neurodevelopment, resulting in long-term behavioral deficits. Therefore, a mouse model that effectively recapitulates these persistent deficits is crucial for furthering our understanding and developing effective interventions.

Given that anti-GluN1 monoclonal NMDAR receptor autoantibodies in human cerebrospinal fluid are sufficient for encephalitis pathogenesis (23), we generated an anti-GluN1 human monoclonal antibody (mAb) from cerebrospinal fluid (CSF) B-cells isolated from a patient with recurrent NMDAR-AE. We then established a novel mouse model of NMDAR-AE by exposing mice to this anti-GluN1 mAb from postnatal day 3 (P3) to P12, a period corresponding to the critical phase in human development from the second trimester to the newborn stage (24). Notably, this age range was selected as it aligns with the onset of a significant portion of pediatric NMDAR-AE cases (25, 26). Remarkably, our mouse model exhibits persistent sensory-motor deficits (19, 27, 28), recapitulating the severe and enduring behavioral deficits commonly seen in NMDAR-AE patients (29, 30). The congruence of these symptoms with human clinical data underscores the relevance of our mouse model for NMDAR-AE, paving the way for a comprehensive investigation into the mechanisms underpinning these persistent deficits.

Our previous study demonstrated that genetic disruption of NMDAR results in callosal projection defects within the primary somatosensory cortex (S1) in mice, underscoring the crucial role of NMDAR signaling in corpus callosum (CC) development (31). The CC, as the largest white matter structure, facilitates connections between various homotopic areas across the two hemispheres of the brain (32). Bilateral sensory-motor coordination relies on the CC bridging the two somatosensory cortices (33, 34). While the general architecture of the CC is established at birth in humans, it continues to develop and mature throughout childhood into adolescence (35–37). Reports show lesions in or volume loss of the CC in NMDAR-AE patients (38–40), with the extent of changes correlating with disease severity (41–43). Consequently, we propose that the inherent vulnerability of the CC to such disruptions might underlie the persistent sensory-motor deficits commonly observed in children with NMDAR-AE.

Therefore, in this study, we employ our novel mouse model to delve into the long-term consequences of transient anti-GluN1 mAb exposure, focusing particularly on morphological changes in callosal axons, their implications for S1 network dynamics, and their subsequent effects on sensory-motor integration during motor behavior. Specifically, in the context of sensory-motor integration, we thoroughly examine functional connectivity during motor tasks, focusing on both inter-hemispheric connections between the left and right primary somatosensory cortices (S1-S1), and intra-hemispheric connections between the primary somatosensory (S1) and primary motor cortex (M1) within the same hemisphere. This comprehensive analysis aims to delineate how alterations at the cellular and network levels contribute to persistent sensory-motor deficits observed in NMDAR-AE.

## Results

### Generation and validation of patient-derived monoclonal anti-NMDAR antibodies

We isolated single, antigen-experienced B-cells from the CSF of a patient with NMDAR-AE using a piezoacoustic liquid handling device (Cellenion, Lyon, France and SCIENION, Berlin, Germany). After RNA extraction and RT-PCR of the heavy and light chains of the immunoglobulin genes of individual cells, we Sanger sequenced the amplicons and produced four distinct patient-derived human IgG1 mAbs using previously published approaches (44, 45) (**Figure 1A**).

**Figure 1:**
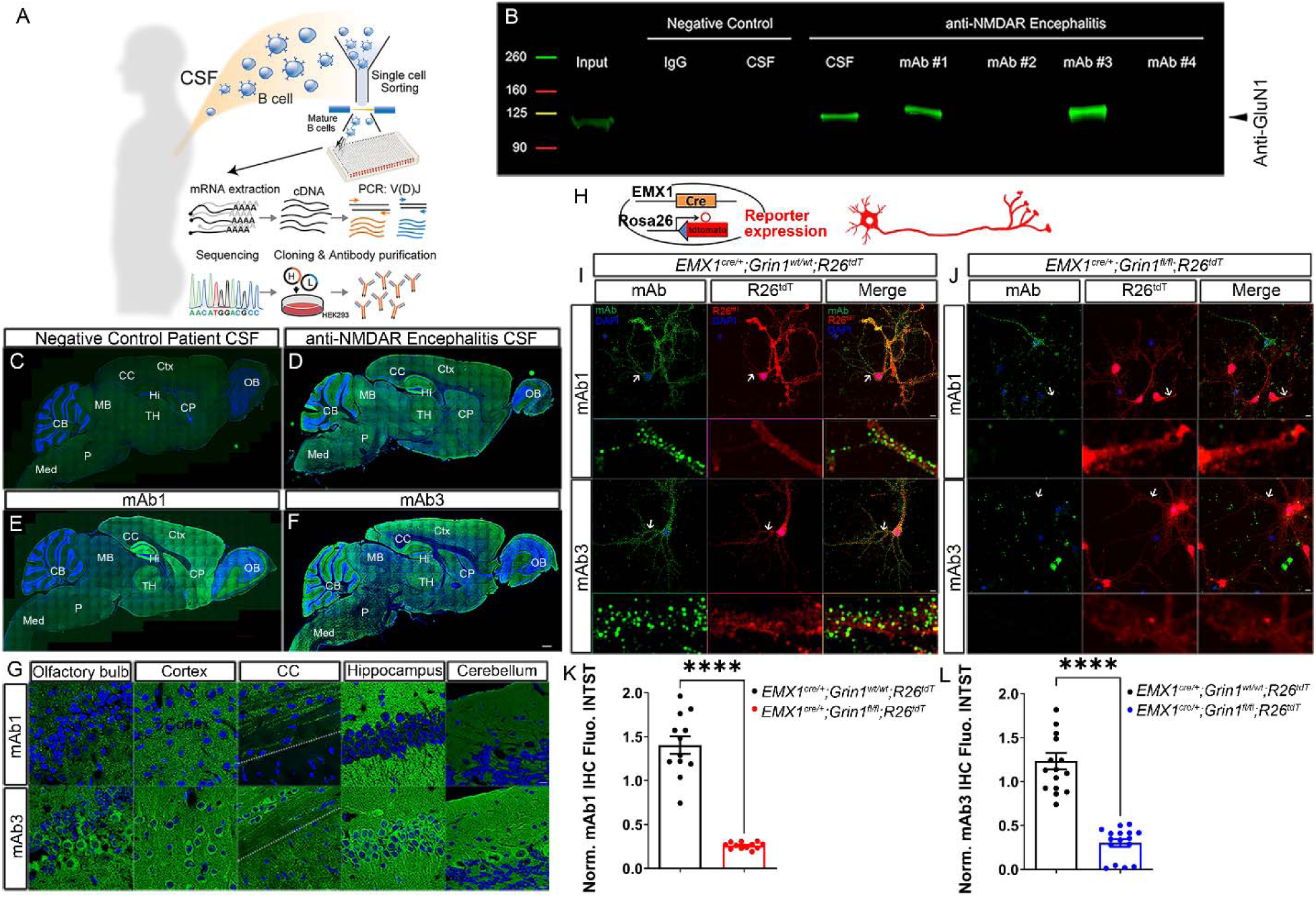
Generation and validation of patient-derived monoclonal anti-NMDAR antibodies. (A) Diagram of generation of patient-derived monoclonal anti-NMDAR antibodies. (B) Western blot demonstrating the immunoprecipitation of GluN1 from postnatal day 40 (P40) mouse brain homogenates using CSF from an anti-NMDAR encephalitis patient, mAb1, and mAb3, which were cloned from the patient’s CSF. Immunoprecipitations with human IgG and CSF from patient without anti-NMDAR encephalitis served as negative controls. Although mAb2 and mAb4 were cloned from the same anti-NMDAR encephalitis patient, they did not immunoprecipitate with GluN1. (C-F) Immunostaining with CSF of negative control patient (C), CSF of anti-NMDAR encephalitis patient (D), mAb1 (E) and mAb3 (F) on sagittal sections of P40 mouse brains. The staining of control patient CSF served as a negative control. (G) Immunostaining pattern of mAb1 and mAb3 across various brain regions. The dash line is the borderline between the cortex and corpus callosum. (H-L) mAb1 and mAb3 recognized extracellular epitopes of NMDAR. (H) We crossed *Emx1^cre/+^*; *Grin1^fl/fl^* mice with Cre-reporter *Rosa26^fs-tdTomato^* mice to produce NMDAR knockout cells labeled with red fluorescence. Hippocampal neurons were cultured from *Emx1^cre/+^*; *Grin1^fl/fl^*; *Rosa26^fs-tdTomato^* mice. Hippocampal cultures of *Emx1^cre/+^*; *Grin1^wt/wt^*; *Rosa26^fs-tdTomato^* mice served as controls. (I, J) mAb1 and mAb3 showed punctate membrane staining in live staining of cultured hippocampal neurons. The staining was gone in red cells of *Emx1^cre/+^*; *Grin1^fl/fl^*; *Rosa26^fs-tdTomato^* cultures (J) but not red cells of *Emx1^cre/+^*; *Grin1^wt/wt^*; *Rosa26^fs-tdTomato^* cultures (I), comfirming mAb specificity for NMDAR. Arrows indicate dendritic fragments, with zoomed-in views provided below each panel. (K, L) Quantification of fluorescence intensity on dendritic fragments shows significant reduction in mAb1 and mAb3 staining in NMDAR knockout neurons compared to controls (****P<0.0001, n = 12 for mAb1, n = 16 for mAb3). Scale bar: 500μm for images C-F; 10μm for images in G, I and J. *R26^tdT^*: *Rosa26^fs-tdTomato^.* Abbreviations: OB, olfactory bulb; Ctx, cortex; Hi, hippocampus; CP, Caudoputamen; TH, thalamus; MB, midbrain; CB, cerebellum; P, pons; Med, medulla.

Immunoprecipitation (IP) with adult mouse brain lysate showed that patient CSF IgG and two of these mAbs (mAb1 and mAb3) bound NMDAR complexes (**Figure 1B**). Compared to the negative control (**Figure 1C**), immunostaining of mouse brain tissue with patient CSF IgG, mAb1, and mAb3 produced intense, characteristic NMDAR neuropil staining in the hippocampus (**Figures 1D-1F**) (2, 9, 46). Immunostaining live murine hippocampal neurons with mAb1 and mAb3 from *Emx1^cre/+^*; *Grin1^wt/wt^*; *Rosa26^fs-tdTomato^* mice, produced punctate cell-surface staining, where red fluorescence marks Cre-mediated recombination without deletion of NMDAR (**Figures 1H, 1I**). The punctate cell-surface staining was lost in red cells of *Emx1^cre/+^*; *Grin1^fl/fl^*; *Rosa26^fs-tdTomato^* cultures (**Figures 1J-1L**), which mark the individual NMDAR knock-out (KO) neurons with red fluorescent protein tdTomato, indicating that mAb1 and mAb3 directly bind NMDAR and recognize extracellular epitopes of NMDAR. This was consistent with previous reports that anti-NMDAR autoantibodies from NMDAR-AE patients recognize the extracellular domain of NMDAR (23, 47, 48).

mAb1 and mAb3 share many of the common features of NMDAR staining. However, mAb3 presents a more uniform pattern (**Figure 1F**), while mAb1 stains stronger in the olfactory bulb, cortex, and hippocampus (**Figure 1E**). Also, mAb3 staining was seen prominently in neuronal cell bodies and processes, whereas mAb1 lacked this pattern (**Figure 1G**). Lastly, mAb1 staining was less prominent than mAb3 staining in postnatal day (P) 8 mouse brains (**Supplementary Figure 1**). These overlapping but different spatial and temporal staining patterns suggested that mAb1 and mAb3 may recognize different subunits of NMDAR.

### Specific binding of mAb3 to the GluN1 subunit significantly decreases NMDAR synaptic currents

NMDARs are heteromeric cation channels with various subunit compositions, characterized by high permeability to Ca²□ ions. To date, seven different subunits have been identified: the GluN1 subunit, four distinct GluN2 subunits (A-D), and two GluN3 subunits. The GluN1 subunit is required for all functional NMDARs (49, 50). In the forebrain, GluN1 primarily assembles with GluN2A and GluN2B to form functional NMDARs. GluN2A-and GluN2B-containing NMDARs follow different developmental expression trajectories with GluN2B as the major GluN2 subunit during the first postnatal week, and GluN2A expression beginning in the first postnatal week and increasing thereafter to become the dominant GluN2 subunit in adults (51–54).

Given the uniform staining pattern of mAb3 during development and in adulthood (**Supplementary Figure 1 and Figures 1F**), we hypothesized that mAb3 may recognize the GluN1 subunit. Supporting this hypothesis, immunoprecipitation-mass spectrometry (IP-MS) analysis indicated that GRIN1 and GRIN2A were the most and second most enriched proteins, respectively (**Supplementary Figure 2**), suggesting mAb3’s strong association with the NMDAR complex. Furthermore, the extensive co-localization of mAb3, but not mAb1, with the anti-GluN1-647 (NR1-647) commercial antibody across dendritic, axonal, presynaptic, and postsynaptic markers, as demonstrated in **Supplementary Figures 3** and 4, further supports the specificity of mAb3 for the GluN1 subunit.

To determine the subunit specificity of mAbs, we analyzed their staining patterns in mice with conditional knockout (cKO) variants of GluN1, GluN2A and GluN2B (31). Specifically, in Grin1cKO (*Emx1^cre/+^*; *Grin1^fl/fl^*; *Rosa26^fs-tdTomato^*) mice, the GluN1 subunit was knocked out, preventing NMDAR assembly in cells marked with red fluorescence. In Grin2a cKO mice (*Emx1^cre/+^*; *Grin2a^fl/fl^)*, GluN2A-containing NMDARs were selectively knocked out in excitatory cortical neurons. Similarly, in Grin2b cKO mice (*Emx1^cre/+^*; *Grin2b^fl/fl^*; *Rosa26^fs-tdTomato^*), GluN2B-containing NMDARs were eliminated in red fluorescence cells. We found the somatic staining of mAb3 was completely absent in red (recombined) cells of Grin1 cKO brain sections (**Figures 2A-2C**), highlighting the dependency of this staining on the presence of the GluN1 subunit. In contrast, the staining of mAb3 was not absent in Grin2a cKO (**Figures 2D-2F**) or Grin2b cKO mice (**Figures 2G-2I**), where GluN2A and GluN2B subunits respectively are knocked out. This implies that mAb3 specifically targets the GluN1 subunit of NMDAR, henceforth referred to as mAb3^[GluN1]^.

**Figure 2:**
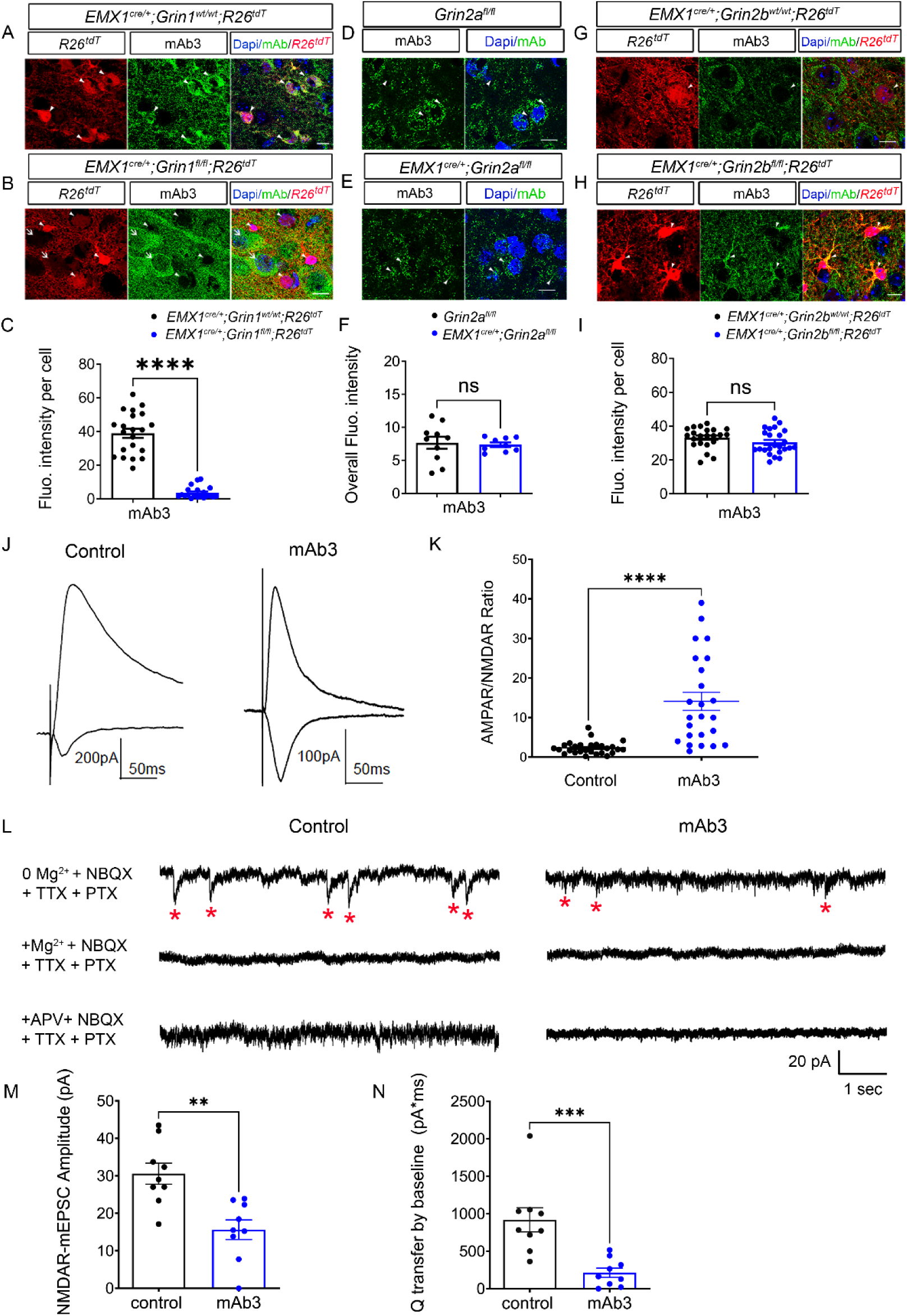
mAb3 specifically binds the GluN1 subunit and significantly decreases NMDAR synaptic currents. (A-I) Characterizing the NMDAR subunit specificity of mAb3 using subunit conditional knockout mice. (A-C) We generated *Emx1^cre/+^*; *Grin1^fl/fl^*; *Rosa26^fs-tdTomato^* mice to produce NMDAR knockout cells labeled with red fluorescence. *Emx1^cre/+^*; *Grin1^wt/wt^*; *Rosa26^fs-tdTomato^* mice of the same litter served as control. Immunostaining of mAb3 was done in these brain sections (A, B). Arrowheads pointed to cells with Cre recombination, while arrow pointed to cells without Cre recombination. mAb3 signals were only detected on non-Cre recombination cells but absent on Cre recombination cells with deletion of NMDAR (B). (C) Quantification of fluorescence intensity of mAb3 immunostaining in single cells (P<0.0001, n = 20 to 21 per group). (D-F) We generated *Emx1^cre/+^*; *Grin2a^fl/fl^* mice to conditional knock out GluN2A-containing NMDAR in excitatory neurons in the cortex. The same littermates *Grin2a^fl/fl^* mice served as controls. There was no difference (F, P=0.79, n = 9 to 10 per group) between the signals of mAb3 in *Grin2a^fl/fl^* mice (D) and *Emx1^cre/+^*; *Grin2a^fl/fl^* mice (E). (G-I) We generated *Emx1^cre/+^*; *Grin2b^fl/fl^*; *Rosa26^fs-tdTomato^*mice to produce GluN2B-containing NMDARs knockout cells labeled with red fluorescence. The littermates *Emx1^cre/+^*; *Grin2b^wt/wt^*; *Rosa26^fs-tdTomato^* mice served as controls. There were no difference between the singals of mAb3 (I, P=0.16, n = 22 to 24 per group) in *Emx1^cre/+^*; *Grin2b^wt/wt^*; *Rosa26^fs-tdTomato^* mice (G) and *Emx1^cre/+^*; *Grin2b^fl/fl^*; *Rosa26^fs-tdTomato^*mice (H). Scale bar: 10μm for all images. (J, K) mAb3 treatment significantly decreases NMDAR synaptic currents in hippocampus slice cultures. (J) Displayed are representative average evoked AMPAR and NMDAR excitatory postsynaptic currents (EPSCs) from slices treated with (mAb3) and without (control) 2 µg mAb3 for 24 hours. NMDAR EPSCs show a significant decrease, and AMPAR/NMDAR ratio shows a significant increase following mAb3 treatment. (K) The left graph represents the AMPAR/NMDAR ratio in mAb3 treated cells (n=24 cells), while the right graph illustrates the ratio in control cells (n=27 cells). Significant differences are indicated by an asterisk (p≤ 0.0001). (L-N) mAb3 blocks NMDAR-mEPSCs in hippocampal neurons. (L) Representative traces of NMDAR-mEPSCs recorded in 0 Mg²[ACSF containing NBQX, TTX, and PTX, followed by addition of 3 mM Mg²[or 50 µM APV to confirm NMDAR-mediated currents. Red asterisks indicate detected NMDAR-mEPSCs. (M) Quantification of NMDAR-mEPSC amplitude in control and mAb3-treated neurons (p<0.01, n=9 cells per group). (N) Charge transfer (Q transfer) by baseline of NMDAR-mEPSCs in control and mAb3-treated neurons (p<0.001, n=9 cells per group). Error bars represent the standard error of the mean (SEM). The above statistics were based on the student T test.

To further investigate the specific effects of mAb3^[GluN1]^, we exposed hippocampal slice cultures to the antibody. After a 24-hour treatment of hippocampal slices with mAb3^[GluN1]^, NMDAR excitatory postsynaptic currents (EPSCs) were significantly decreased, and the AMPAR/NMDAR ratio was significantly increased (**Figures 2J and 2K**). Additionally, recordings of NMDAR-mediated miniature excitatory postsynaptic currents (NMDAR-mEPSCs) further confirmed the inhibitory effects of mAb3^[GluN1]^. Specifically, mAb3^[GluN1]^ treatment significantly reduced the amplitude of NMDAR-mEPSCs (**Figure 2M**) and resulted in a significant decrease in charge transfer by baseline (**Figure 2N**). The specificity of these NMDAR-mEPSCs was confirmed by their sensitivity to 3 mM Mg²[or 50 µM APV, both of which abolished the currents (**Figure 2L**). These findings support our inference that mAb3^[GluN1]^ selectively binds to the GluN1 subunit, thereby inhibiting NMDAR function. According to Kreye et al. (2016) (23), autoantibodies targeting the GluN1 subunit of NMDAR are believed to be the main autoantibodies responsible for NMDAR-AE. Therefore, in this study, we chose to focus our investigation on the effects of mAb3^[GluN1]^ to explore the underlying mechanisms behind persistent sensory-motor deficits in a mouse model.

### mAb3^[GluN1]^ injection disrupts callosal projection patterns in primary somatosensory cortex

Persistent sensory-motor deficits are common in pediatric NMDAR-AE patients (19, 27, 28). Our previous work demonstrated the vulnerability of the primary somatosensory cortex (S1) callosal circuit, critical for bilateral sensory-motor coordination, to disruption by commercial rabbit-derived anti-NMDAR antibodies when injected from P3 to P12 in mice (31). This developmental window is crucial for the formation and maturation of callosal projections in mice and corresponds to the critical phase in human development from the second trimester to the newborn stage (24).

The period from P3 to P12 encompasses key developmental stages of S1 callosal projections. During this time, S1 callosal axons transition from one hemisphere to the other (P3), reach the white matter beneath the contralateral S1 (P5), spread within the contralateral S1 (P8), and finally undergo a pruning process to mature into a network confined to the border between the primary and secondary somatosensory cortices (S1/S2) (P14). We now turned our attention to whether the patient-derived anti-NMDAR autoantibody mAb3^[GluN1]^ might have a similar impact on the development of the S1 callosal circuit. To address this question, we used *in utero* electroporation to label layer II/III callosal neurons with enhanced green fluorescent protein (EGFP). This procedure was performed on progenitor cells at embryonic day (E) 15.5 (**Figure 3A**). We then administered mAb3^[GluN1]^ (1.6 μg) into the lateral ventricular zone of the contralateral target S1 of mice twice daily from P3 to P12. Our previous study (31) demonstrated the feasibility and efficiency of this cortical antibody injection (Supplementary Figure 5).

**Figure 3:**
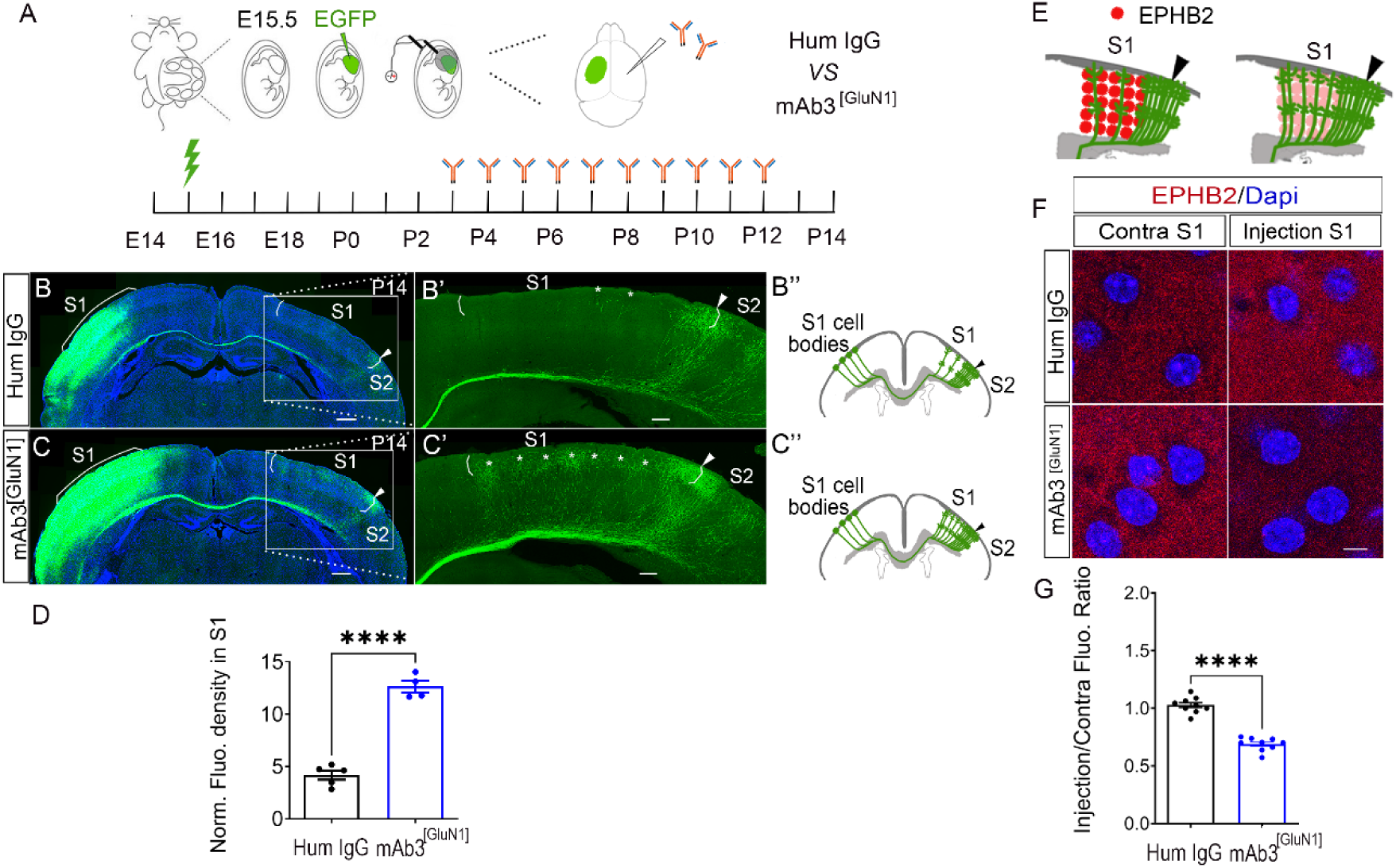
Disrupted callosal projections in primary somatosensory cortex (S1) after intraventricular injection of mAb3^[GluN1]^ from P3 to P12. (A) Diagram of the experimental procedure. EGFP plasmid was injected into the lateral ventricle of the embryo at embryonic day15.5 (E15.5) and an electrical pulse was given to enable the plasmid to enter cortical progenitor cells of layer II/III in the ventricular zone. mAb3^[GluN1]^ was injected into the lateral ventricle from P3 to P12 in contralateral cortex. Human IgG served as control. Compared with control (B), mAb3^[GluN1]^ injection mice showed dramatically increased callosal projections (C) in S1 at P14. “*” pointed to the callosal axons in S1. (D) Quantification of the fluorescence density. Human IgG *VS* mAb3^[GluN1]^: P < 0.0001. n = 4 to 5 per group. (E) Diagram of EPHB2 expression in S1. (F) Expression of EPHB2 in S1 of injecting side and contralateral noninjecting side for the two treatments. (G) Quantification of fluorescence intensity ratio of injecting side to contralateral noninjecting side. Human IgG *VS* mAb3^[GluN1]^: P < 0.0001. n = 9 for each group. Scale bar: 500μm for B, B’, C, C’; 5μm for F. Above statistics were based on Mann-Whitney test.

Upon examining the S1 callosal projection pattern at P14, we observed a typical, orderly neural network in control human IgG-treated mice (human IgG, 1.6 μg, Jackson ImmunoResearch). The callosal projections were primarily localized around the S1/S2 border, with a comparatively sparse distribution in S1 (**Figure 3B**). However, in the mAb3^[GluN1]^-treated mice, the callosal projections in S1 were dispersed and significantly increased in number, leading to a complete alteration of the normal organization of projections within this region (**Figure 3C and 3D**, P < 0.0001). These findings strongly suggest that mAb3^[GluN1]^ injections, when administered during this critical period of S1 callosal circuit development, markedly interfere with the normal formation of these neural networks.

Interestingly, NMDAR, a well-established neurotransmitter receptor, seems to play a role in neural circuit formation. Our previous genetic study with Grin1 knockout model demonstrated similar circuit disruptions (31). We showed that these changes are not mediated by NMDAR’s ion channel function, but by interactions with the axon guidance pathway EPHRIN-B/EPHB, particularly EPHRIN-B1 and EPHB2. The loss of NMDAR resulted in a selective reduction of EPHB2 in the knockout cells. As EPHRIN-B1-EPHB2 is a repulsive guidance cue, this loss resulted in an increase in callosal projections into S1.

In this current study, we observed a significant reduction of EPHB2 expression in the hemisphere injected with mAb3^[GluN1]^ (**Figure 3E-3G**). This reduction not only substantiates our hypothesis that mAb3^[GluN1]^ downregulates the EPHRIN/EPH pathway, leading to an increase in callosal projections, but also underscores the lasting impact that temporary disruption of NMDAR function by mAb3^[GluN1]^ during a critical developmental window can have on neural circuitry.

### Persistent sensory-motor deficits in mice following transient developmental exposure to mAb3^[GluN1]^

Given our findings of S1 callosal circuit disruption, we sought to investigate whether these neurodevelopmental changes could be reflected in the long-term behavior of mAb3^[GluN1]^-treated mice. Drawing parallels with the clinical presentations observed in pediatric patients, we hypothesized that this transient developmental exposure to the antibody could result in long-lasting sensory-motor deficits. Accordingly, we conducted a series of behavioral tests on these mice when they were between 1 to 4 months, each test designed to probe different aspects of sensory-motor integration.

To ensure the validity of our sensory-motor assessments, we first performed control tests—the elevated plus maze (EPM) test and the open field test (OFT)—to rule out potential confounding factors such as anxiety and overall changes in locomotor activity. In both tests, no significant differences were observed between the groups (**Supplementary Figure 6 f**or EPM; **Figures 4A and 4B** for OFT), confirming that the performance in our behavioral tasks was not confounded by differences in anxiety levels or overall locomotor activity.

**Figure 4:**
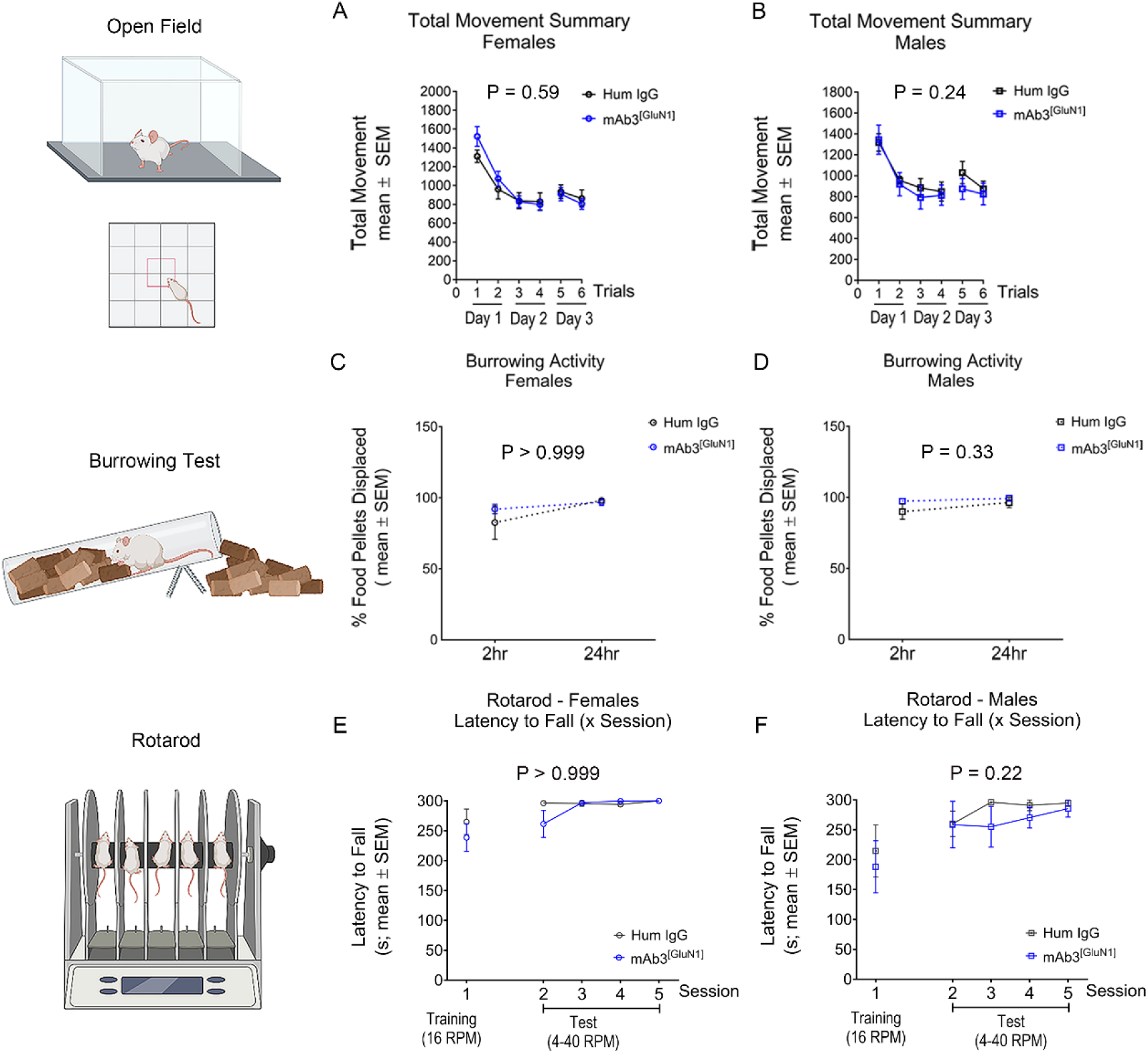
Unaffected gross movements in mAb3^[GluN1]^-treated mice. (A, B) Locomotor activity in Open Field Test (OFT). There was no difference in total movements between Human IgG and mAb3^[GluN1]^-treated female (A) and male (B) mice. (C, D) Digging and kicking activity in burrowing test. There was no difference in burrowing performance between the two groups in females (C) and males (D) mice. (E, F) Motor coordination in rotarod. There was no difference in rotarod performance between the two groups in females (E) and males (F) mice. n = 6 to 8 per group. P value on each figure graph represents the statistical difference between the two groups over trials by using Two-way Analysis of Variance (ANOVA) and Geisser-GreenHouse correction.

To evaluate gross motor skills and coordination, we employed the burrowing test and the rotarod test. The former focuses on digging and kicking activity, while the latter evaluates gross motor coordination. Performance in these tasks (**Figures 4C-4F**) was comparable between mAb3^[GluN1]^-treated mice and the control groups.

While the mAb3^[GluN1]^-treated mice demonstrated similar abilities in gross motor skills and coordination tasks, their performance was significantly impaired in tasks requiring fine motor skills and complex sensory-motor coordination. These included nest building (**Figure 5B**), balance beam walking (**Figure 5D**, **Video S2**), and the facing upward pole test (**Figures 5L and 5M**, **Videos S5-S8**)—all demanding sophisticated coordination and intricate sensory-motor integration. Notably, male mAb3^[GluN1]^-treated mice showed more severe impairments than their female counterparts (**Table 1**).

**Figure 5:**
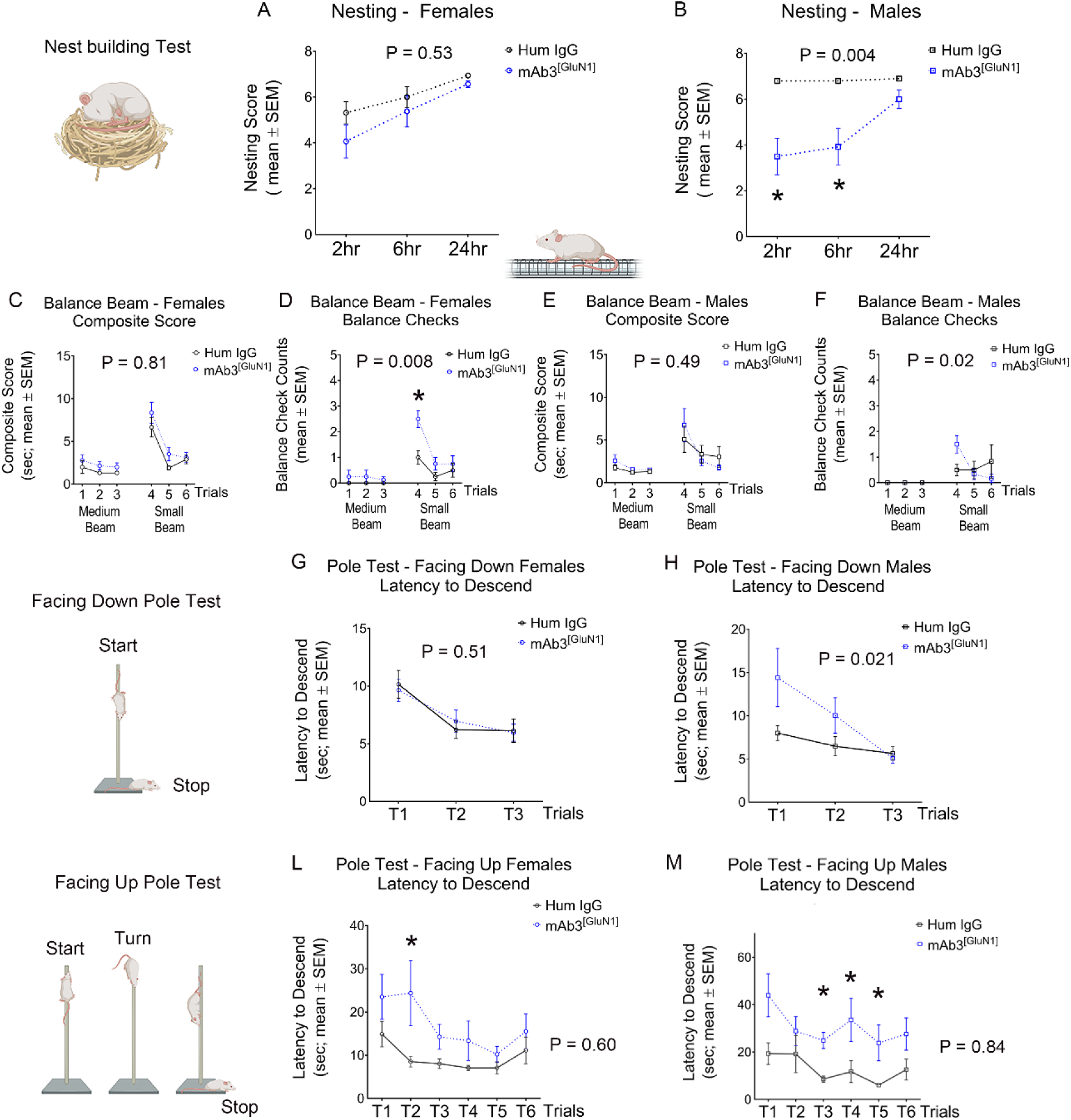
Persistent impaired fine movements in mAb3^[GluN1]^-treated mice. (A, B) Nest building. There was significantly impaired nest building at 2hr (Human IgG *VS* mAb3^[GluN1]^: P = 0.02) and 6hr (Human IgG *VS* mAb3^[GluN1]^: P = 0.03) in mAb3^[GluN1]^-treated male (B) mice. (C-F) Balance beam. There was no difference in latency of beam crossing for female (C) and male (E) mice. However, there was a significantly increased balance check in mAb3^[GluN1]^-treated female mice (D) at Trial 4 (Human IgG *VS* mAb3^[GluN1]^: P = 0.009). (G, H) Facing down pole test. A significant difference was observed between mAb3^[GluN1]^-treated and Human IgG-treated male mice over all three trials (H). However, there was no difference in descent latency in each trial for female (G) and male (H) mice. (L, M) Facing upward pole test. There was significantly increased descent latency in mAb3^[GluN1]^-treated female mice (L) at Trial 2 (Human IgG *VS* mAb3^[GluN1]^: P = 0.01), and mAb3^[GluN1]^-treated male (M) at Trial 3 (Human IgG *VS* mAb3^[GluN1]^: P = 0.04), Trial 4 (Human IgG *VS* mAb3^[GluN1]^: P = 0.04), and Trial 5 (Human IgG *VS* mAb3^[GluN1]^: P = 0.03). n = 6 to 8 per group. P value on each figure graph represents the statistical difference between the two groups over all trials by using Two-way ANOVA and Geisser-GreenHouse correction. The p value in figure legends represents the statistical difference between two treatments for one trial by using Dunn’s multiple comparisons test.

**Table 1.** Summary of impaired fine movements in mAb3^[GluN1]^-treated mice. Locomotor activity, muscle strength, and muscle coordination served as controls for baseline movement activity.

| Treatment | Gender | Locomotor Activity | Muscle Strength | Muscle Coordination | Nest Building | Balance Check | Latency of Facing Up Pole |
| --- | --- | --- | --- | --- | --- | --- | --- |
| mAb3<sup>[GluN1]</sup> | **Female** | – | – | – | – | ↓ | ↓ |
| mAb3<sup>[GluN1]</sup> | **Male** | – | – | – | ✕ | – | ↓ |

–: No Change; ↑: Increased; X: Impaired

It is important to note that no correlation was observed between motor performance and body weight, muscular strength, or coordination in either the mAb3^[GluN1]^-treated or control mice, as shown in **Supplementary Figures 7-10**. Intriguingly, male mAb3^[GluN1]^-treated mice exhibited significantly better muscle coordination than control mice (**Supplementary Figure 10D**), a finding that inversely correlates with their performance deficits in the challenging sensory-motor task, the facing upward pole test (**Figure 5M**). This suggests that the observed sensory-motor deficits are primarily due to neural circuitry disruptions caused by the mAb3^[GluN1]^ exposure. These results demonstrate that transient exposure to pathogenic anti-NMDAR autoantibodies can lead to persistent fine motor coordination deficits in mice.

### Disrupted inter-hemispheric functional connectivity in S1 of mAb3^[GluN1]^-treated mice

Considering the persistent sensory-motor deficits observed in mAb3^[GluN1]^-treated mice, particularly in fine movements, we hypothesized that disruptions in sensory-motor integration might underlie these deficits. Functional connectivity, defined as the statistical dependency or synchrony of neuronal activity across different brain regions, is a key determinant of sensory-motor integration (55, 56). Alterations in functional connectivity within relevant cortices can disrupt this integration, manifesting as behavioral deficits.

As depicted in **Supplementary Figure 11**, inter-hemispheric connections between primary somatosensory cortices (S1s) are crucial for bilateral sensory integration, allowing for coordinated sensory-motor processing across hemispheres (57). In contrast, intra-hemispheric connections between S1 and the primary motor cortex (M1) are key for linking sensory input to motor output within the same hemisphere, essential for executing motor commands (55, 58, 59). Therefore, to understand whether alterations in this functional connectivity might underlie the observed behavioral deficits, we placed a 30-channel multi-electrode array (MEA) electroencephalogram (EEG) spanning the mouse skull (60) (**Figure 6A**). This array allowed us to conduct simultaneous recordings from various cortical regions across both hemispheres, including S1 and M1, during the performance of sensory-motor tasks by mAb3^[GluN1]^-treated male mice. Male mice were specifically selected for this analysis due to their more severe deficits in sensory-motor coordination task performance.

**Figure 6:**
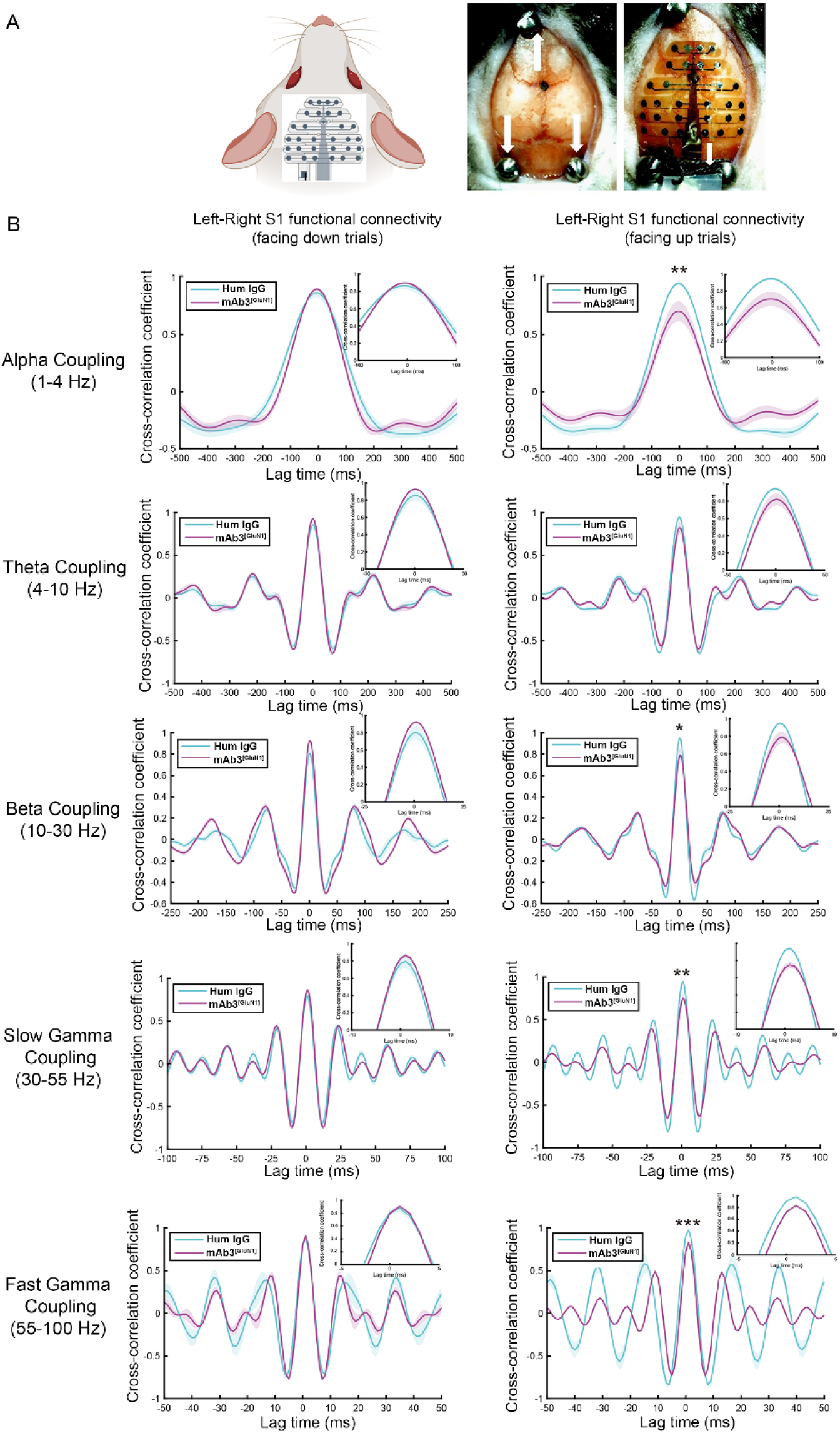
Disrupted inter-hemispheric functional connectivity in S1 of mAb3^[GluN1]^-treated male mice. (A) Schematic representation of the 30-channel EEG array and the process of its implantation on the mouse skull (Adapted from Jonak et al., 2018). As previously described, mAb3^[GluN1]^ was injected into the right hemisphere of mice from P3 to P12, with EEG surgery and recording carried out when the mice were between 2-3 months old. We recorded EEG signals while mice performed the facing down and facing upward pole tests. (B) The cross-correlation coefficient curve for left-right S1 functional connectivity during the pole test in both ’facing down’ and ’facing up’ trials. In the graph, Hum IgG-treated male mice (n=3) served as control for mAb3^[GluN1]^-treated male mice (n=3). The control group demonstrated significantly higher left-right S1 functional connectivity compared to the antibody group during ’facing up’ trials. This difference was not observed during ’facing down’ trials. The frequency bands where differences were observed include alpha, beta, slow gamma, and fast gamma. Differences in functional connectivity between treatment groups and conditions were assessed using two-way ANOVA followed by Šídák’s multiple comparisons test. Data are presented as mean ± SEM. Statistical significance was set at p < 0.05. The waveform in the figure was plotted using MATLAB, with shaded areas representing SEM.

This was substantiated during the challenging facing upward pole test, which revealed significant behavioral deficits (**Figure 5M**). In this task, we observed a significant reduction in inter-hemispheric functional connectivity between the left and right S1 across multiple frequency bands (**Figure 6B** and **Supplementary Figure 13A**) in mAb3^[GluN1]^-treated male mice. In contrast, such a reduction was absent in the less demanding facing down pole test (**Figure 6B**), where no behavioral deficits were observed (**Figure 5H**).

Importantly, we did not identify any significant alteration in intra-hemispheric functional connectivity between S1 and M1 within the right hemisphere, where the antibody was injected, during either task (**Supplementary Figure 13B**). These findings suggest that the deficits induced by mAb3^[GluN1]^ primarily affect inter-hemispheric connections, sparing intra-hemispheric connections. Given the crucial role of S1 callosal axons – the structural basis for inter-hemispheric connections (57, 59, 61)– we next turned our investigation to potential permanent morphological alterations in these axons in adult mice.

### Permanent alterations of S1 callosal axons in mice exposed transiently to mAb3^[GluN1]^ during development

To build upon our functional connectivity findings, we next sought to identify morphological changes that might correlate with the alterations in the S1 callosal circuit. We examined the S1 callosal circuit at 4 months, after mice had performed all behavioral tasks. Interestingly, by 4 months of age, the excess S1 callosal projections seen in mice treated with mAb3^[GluN1]^ during development had resolved (**Supplementary Figures 14A** and 14B). However, we also took a closer look at the cellular level, tracing individual callosal axon terminals in S1 (**Figures 7A**, **Supplementary Figure 15, Methods**) and assessing various morphometric features.

**Figure 7:**
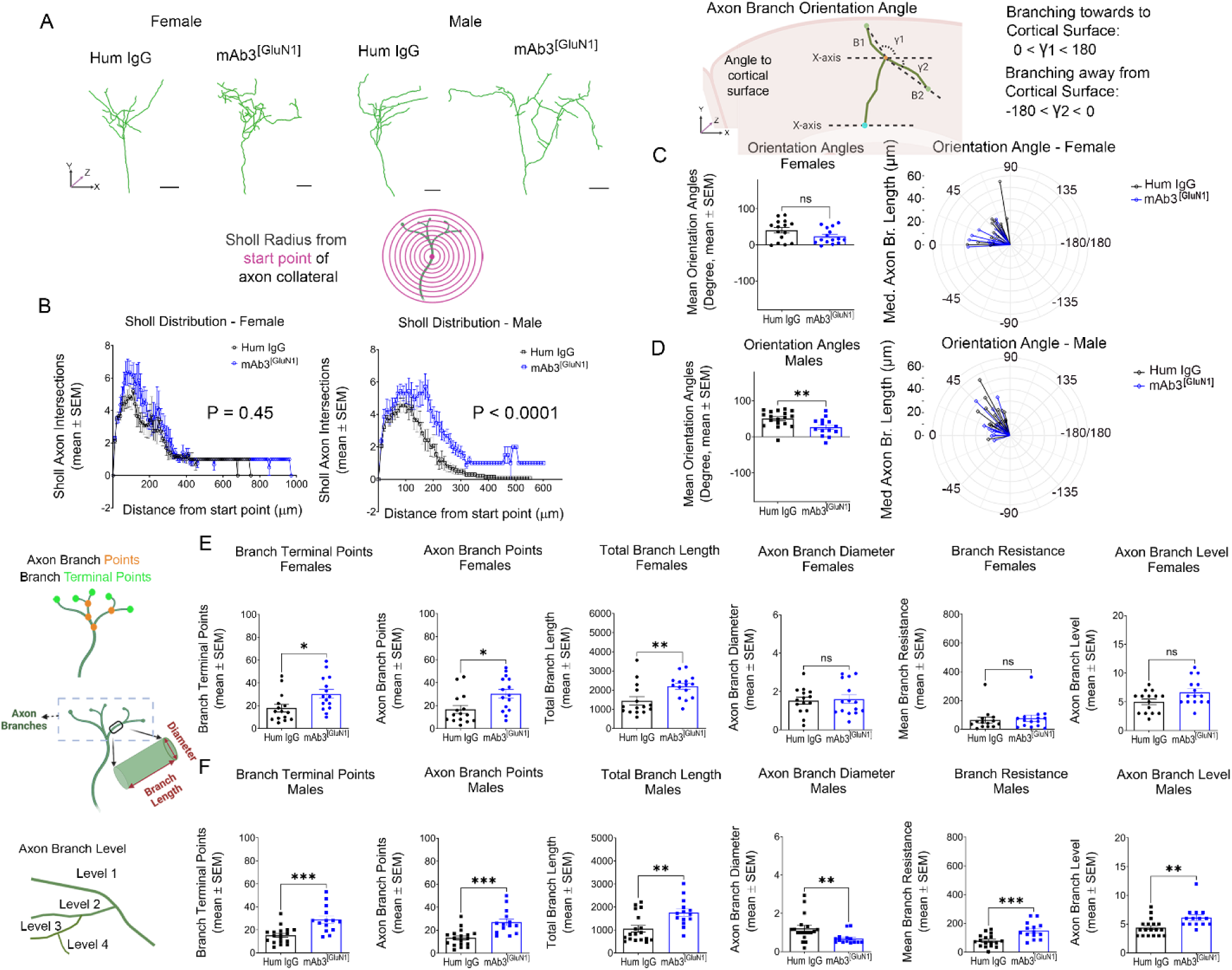
Permanent morphological alterations of S1 callosal axons in mAb3^[GluN1]^-treated mice. (A) Represented morphology of individual callosal axon terminals in S1 at 4 months. (B) Sholl analysis of callosal axon branches in female and male mice. Concentric circles from the start point are used to count the number of axon intersections. Start point was defined at 10μm before the first branch intersection along the main axon trunk (magenta point in the diagram). Axon branch orientation angle in female (C) and male (D) mice. The angle forms between the extending line connecting the distal axon branch segment to the x-axis of each image within the XY plane. X-axis is parallel to the cortical surface. The angle is from -180 to 180 degrees and is used to quantify the extending direction of the axon branch to the cortical surface. A positive angle (0 -180) means the axon branch is extending towards to cortical surface, while a negative angle (-180 -0) means the axon branch is extending away from the cortical surface. (E, F) Morphological features of axon branch terminals in mAb3^[GluN1]^-treated female (E) and male (F) mice. \**P* < 0.05; *\**\**P* < 0.05; \*\*\**P* < 0.01. n = 14 to 18 per group (B-F). For the Hum IgG-treated group, 5 female mice (15 terminals) and 4 male mice (18 terminals) were analyzed. For the mAb3^[GluN1]^-treated group, 4 female mice (14 terminals) and 6 male mice (14 terminals) were analyzed. Each mouse had 3-4 terminals analyzed. The above statistics were based on Mann-Whitney test. The plots in C and D were made in R using ggplot2 package.

Remarkably, we found that the morphology of axon terminals in S1 was permanently altered in a sex-specific manner in mAb3^[GluN1]^-treated mice, even months after the transient exposure to the antibody (**Table 2**). The male mice displayed significantly higher numbers of axon branch crossings, suggesting increased branch complexity and larger terminal field areas (62). This conclusion was drawn based on the results of Sholl analysis, a method used to quantify the number of intersections of axon terminals with concentric circles (**Figure 7B**). Both mAb3^[GluN1]^-treated female and male mice displayed significantly increased axon branch points, terminal points, and total branch length (**Figures 7E, 7F**), further indicating higher branch complexity. Male mice also showed a reduced branch diameter, suggesting increased axon resistance, which could potentially decrease signal conduction velocity (**Figure 7F**). Changes in axon branch levels and angles were also observed (**Figure 7F**, **Supplementary Figure 16**) in male mice, along with alterations in branch orientation angles (**Figures 7C, 7D**).

**Table 2.**
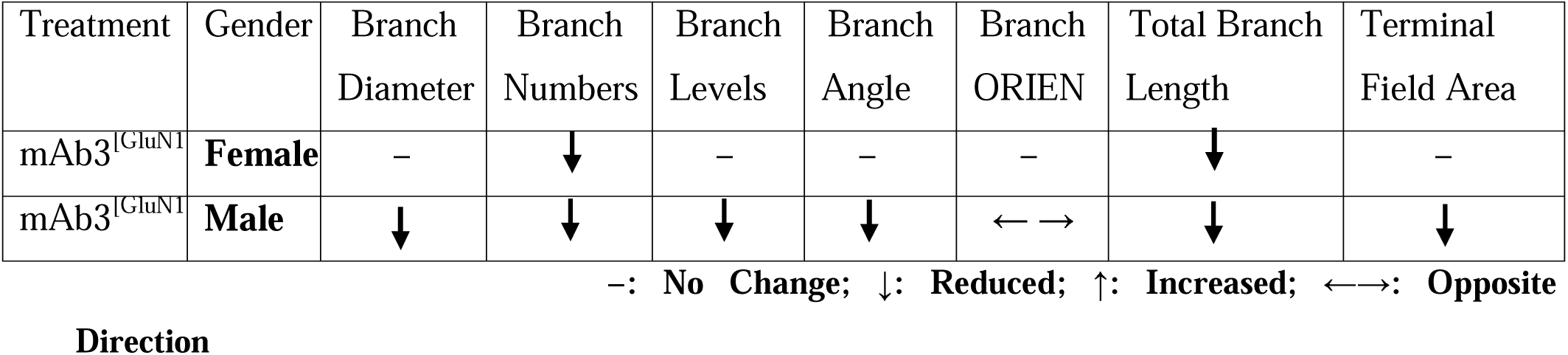
Summary of morphological alterations in S1 callosal axons for mAb3^[GluN1]^-treated mice.

| Treatment | Gender | Branch Diameter | Branch Numbers | Branch Levels | Branch Angle | Branch ORIEN | Total Branch Length | Terminal Field Area |
| --- | --- | --- | --- | --- | --- | --- | --- | --- |
| mAb3<sup>[GluN1]</sup> | **Female** | – | ↓ | – | – | – | ↓ | – |
| mAb3<sup>[GluN1]</sup> | **Male** | ↓ | ↓ | ↓ | ↓ | ← → | ↓ | ↓ |

–: No Change; ↓: Reduced; ↑: Increased; ←→: Opposite

Strikingly, these morphological alterations of S1 callosal axons showed a positive correlation with the behavioral deficits in sensory-motor coordination. The mAb3^[GluN1]^-treated male mice, who displayed the worst sensory-motor coordination performance (**Table 1**), also exhibited the most significantly altered morphology in S1 callosal terminals (**Table 2**). This correlation suggests that callosal termination defects may indeed underlie the long-lasting sensory-motor deficits seen in our mouse model of NMDAR-AE.

### Aberrant network excitability in S1 in mice exposed transiently to mAb3^[GluN1]^ during development

Building on these findings, we next examined whether these observed morphological alterations in S1 callosal axons were accompanied by changes in network excitability. We thus performed ex vivo electrophysiological recordings of S1 while stimulating the CC underneath. This approach aimed to assess the potential effects of any structural changes in CC axons projecting into the S1 cortex. These recordings were conducted on acute brain slices from male mice at 6 months of age, following transient exposure to mAb3^[GluN1]^ during development. The evoked cortical responses were recorded across all the layers of the S1 cortex using a linear 16-channel electrode array (**Figure 8A**, **Methods**).

**Figure 8.**
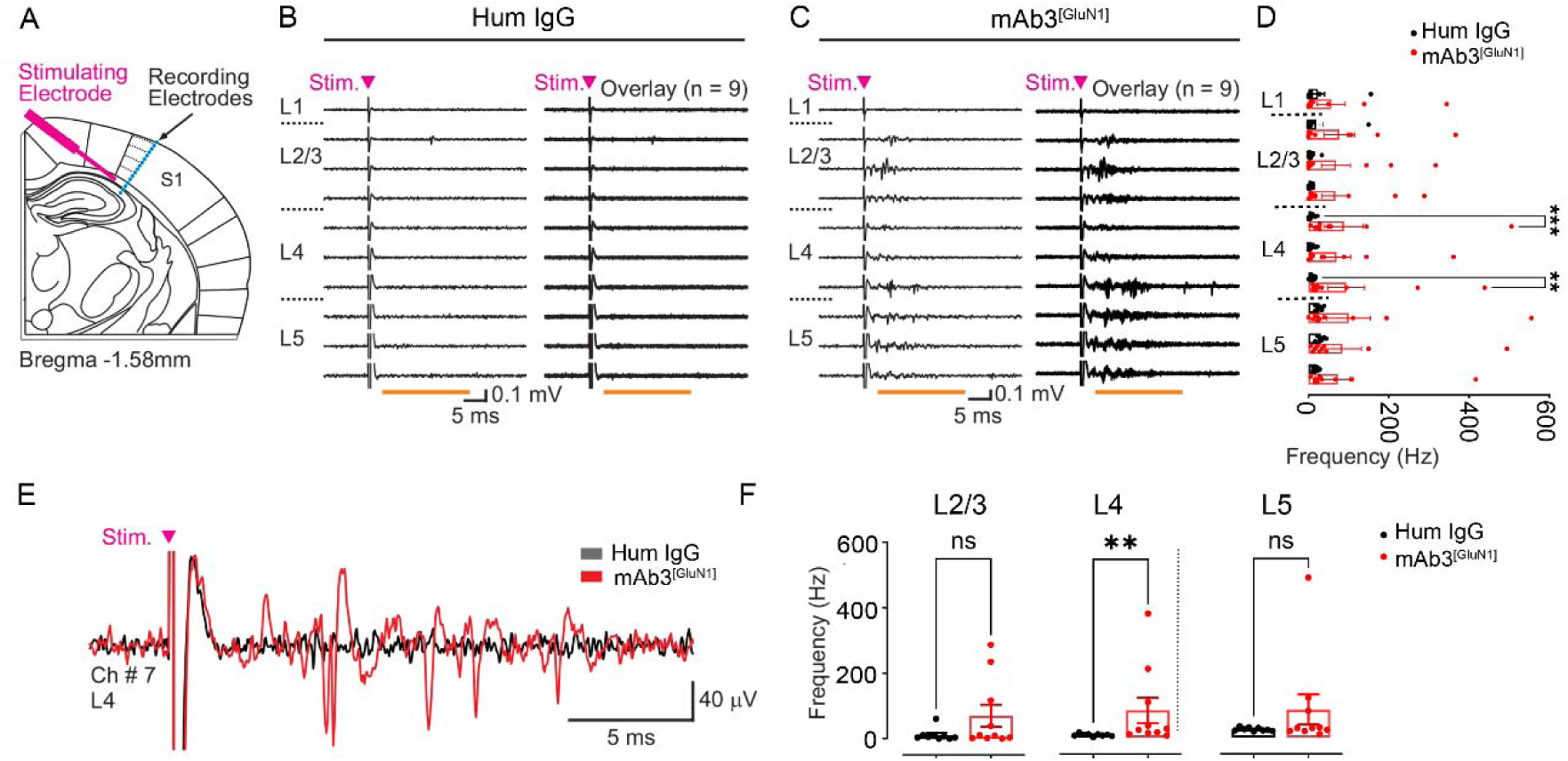
The primary somatosensory cortex is hyperexcitable in mAb3^[GluN1]^-treated male mice at 6 months. (A) Schematic of the experimental design of ex vivo recordings showing the location of the stimulating electrode in the white matter and the extracellular recording array spanning all the layers in the S1 cortex (blue). (B, C) Ex vivo recordings of putative extracellular spikes in response to a 500 μA electrical pulse stimulation (arrowhead) from a brain slice of a control Hum IgG-treated mouse (B) and a brain slice from mAb3^[GluN1]^-treated mouse (C). In each case, a single trace and an overlay of 9 traces are presented from the same slice. Evoked spikes were counted during the time window indicated by the horizontal orange line. (D) Mean frequency of spikes for channels located in layers 1-5. Data: mean ± SEM; n = 9 slices from 4 Hum IgG-treated mice and n = 10 slices from 4 mAb3^[GluN1]^-treated mice. P values are from the Mann-Whitney Rank Sum test comparing Hum IgG vs mAb3^[GluN1]^ for each channel. Note that only channels 5 and 7 (both located in layer 4) show significant differences between control and mAb3^[GluN1]^ groups. Using Kruskal-Wallis ANOVA with multiple comparisons (Dunn’s method) with 19 degrees of freedom p<0.001 between groups. (E) Traces from the slices in (B, C) showing evoked spikes from channel 7 located in layer 4 (L4). (F) Same quantification as in E but with channels grouped for layers 2/3 (averaged across channels 2–4), layer 4 (averaged across channels 5– 7), and layer 5 (averaged across channels 8–10). Note that although layers 2/3 and 5 show tendency for higher spiking rate in Hum IgG vs mAb3^[GluN1]^ groups, only layer 4 shows a significant different between groups. Data: mean ± SEM; n = 9 slices from 4 control Hum IgG-treated mice and n = 10 slices from 4 mAb3^[GluN1]^-treated mice. P value is from the Mann-Whitney Rank Sum test comparing Hum IgG vs mAb3^[GluN1]^ for each channel. Kruskal-Wallis ANOVA with multiple comparisons (Dunn’s method) with 5 degrees of freedom: p<0.001 and using this method only layer 4 shows significant difference between Hum IgG and mAb3^[GluN1]^ groups (p<0.05 with multiple comparisons, 5 degrees of freedom).

We found that CC stimulation resulted in significantly higher evoked spiking in layer 4 in the mAb3^[GluN1]^-treated mice compared to controls (**Figures 8B-8F**). These findings are consistent with our earlier observations of altered inter-hemispheric functional connectivity, particularly between S1 regions (**Figure 6B**), suggesting that structural changes in callosal axons may impact S1 network excitability. Together, these results suggest a link between structural alterations in callosal axons, changes in network excitability, and disrupted sensory-motor integration in mAb3^[GluN1]^-treated mice, contributing to their persistent sensory-motor deficits.

## Discussion

Substantial progress has been made in understanding the acute phase pathophysiology of NMDAR-AE (63). However, psychomotor abnormalities, particularly sensory-motor coordination impairments, often persist even after the autoantibody titers have declined, especially in pediatric patients (64). These underlying mechanisms of persistent impairments have remained largely unexplored. To address this, we successfully developed a novel mouse model that recapitulates some of the long-lasting sensory-motor deficits seen in NMDAR-AE patients. This model was established by administering a patient-derived GluN1-specific mAb into the lateral ventricular zone of neonatal mice during a period known to be essential for callosal projection development. Using this model, we then investigated the cellular and circuit mechanisms underlying these persistent sensory-motor coordination impairments.

### Morphological Changes in Callosal Axons

Our data demonstrate that transient exposure of mAb3^[GluN1]^ from P3 to P12 — a period corresponding to the critical phase in human development from the second trimester to the newborn stage (24) — causes significantly increased S1 callosal projection at P14 (**Figure 3**) and permanent morphological alterations in S1 callosal axons, characterized by increased axon branch complexity and altered branch orientation angles (**Figure 7 and Supplementary Figure 16**).

Hand and finger coordination rely heavily on communication through the CC in the somatosensory cortex (33, 34). Sensory information from the two halves of the body is integrated bilaterally at the cortical level via the CC (34). Clinical studies suggest that the extent of axonal tract pathology in NMDAR-AE patients correlates with their disease severity (41–43). Notably, volume loss of the CC has been observed in a 2-year-old child with a delayed diagnosis of NMDAR-AE, who presented with severe symptoms and persistent neurological impairment (39). Our data and clinical observations highlight the likely involvement of the CC in brain network, such as sensory-motor brain network alterations in NMDAR-AE.

### Network and Functional Connectivity Alteration

These morphological changes in callosal axons were associated with an aberrant recruitment of S1 cortex by CC stimulation, as shown by our ex vivo electrophysiological findings from mAb3^[GluN1]^-treated mice. Specifically, electrical stimulation of CC resulted in strong increase in evoked spiking in layer 4 of S1 (**Figure 8**), consistent with the increased number of CC axonal arborizations in S1 (**Figure 7**). This unilateral increase in S1 network excitability could disrupt the precise timing needed for effective sensory-motor integration and coordination (65), potentially contributing to the coordination deficits observed in mAb3^[GluN1]^-treated male mice during complex sensory-motor tasks, such as nest building and the facing upward pole test (**Figures 5B, 5M**).

Our MEA EEG data further revealed a significant reduction in inter-hemispheric functional connectivity between S1 regions during the challenging facing upward pole task, but not in the less demanding facing downward pole task (**Figure 6 and Supplementary Figure 13**). This indicates that the S1-S1 callosal structural deficits specifically affect fine sensory-motor movements that require multiple feedback loops (**Supplementary Figure 11**), where accumulated imprecise timing becomes more apparent. No changes were detected in intra-hemispheric connectivity between S1 and M1 (**Supplementary Figure 13**), underscoring the specific role of S1-S1 connectivity in complex sensory-motor coordination. The pattern suggests that the morphological changes in S1 callosal axons—essential for inter-hemispheric communication—are critical contributors to sensory-motor impairments.

While our ex vivo electrophysiological recordings suggest that postsynaptic changes result from increased network excitability, we have not directly assessed the potential contributions of dendritic complexity in S1 neurons. Therefore, we cannot rule out the possibility that mAb3^[GluN1]^ effects on dendrites or other brain regions may also contribute to the observed changes. Future studies will explore these mechanisms, including why hyperexcitability is more prominent in layer 4 and whether mAb3^[GluN1]^ affects thalamocortical pathway or intrinsic neuronal properties, and their impact on sensory-motor integration deficits.

Our findings suggest that transient exposure to the GluN1 mAb can induce irreversible morphological changes in callosal axons, disrupting sensory-motor integration and resulting in persistent sensory-motor deficits. This study provides insights into cellular and network-level mechanisms underlying long-term sensory-motor deficits in NMDAR-AE. Although our results are based on a mouse model, they hint at a plausible scenario in young children, where exposure to anti-NMDAR autoantibodies during a critical developmental window could lead to similar morphological alterations in S1 callosal neurons, with potential long-term consequences for sensory-motor integration and coordination.

Furthermore, a recent functional magnetic resonance imaging (fMRI) study reported significant reductions of functional connectivity in the sensory-motor network in NMDAR-AE patients (43), strongly supporting our cellular and circuitry findings. These consistent discoveries underscore the validity of our model and hint that the mechanisms identified in this study might contribute to the long-term deficits seen in NMDAR-AE patients. Further investigations are needed to confirm whether similar morphological changes and disruptions in functional connectivity occur in humans with NMDAR-AE and to determine if these changes correlate with the severity and persistence of behavioral deficits.

### Sex Differences in Response to mAb3^[GluN1]^

The administration of the anti-GluN1 mAb from postnatal days 3 to 12 coincides with a critical period for estrogen’s influence on sex differentiation in developing mice. Estrogen receptor expression, particularly estrogen receptor alpha (ERα), surges in female mice around this time, overlapping with crucial stages of synaptogenesis and neural circuit formation (66). This observation raises the possibility that estrogen could modulate the neural response to mAb exposure, potentially explaining the sex differences observed in our study. Estrogens are known to influence glial cell functions and synaptic configuration—both pivotal for neural circuit development and resilience to injuries (67, 68). By activating estrogen receptors, estrogens not only enhance the maturation and survival of neurons but also promote synaptic plasticity and resistance to inflammatory damage. The differential impact of estrogens during the antibody exposure window could therefore lead to the distinct morphological and behavioral outcomes noted between sexes (**Tables 1 and 2**). Understanding the mechanisms behind these sex differences may inform treatment strategies for NMDAR-AE and highlights the need for further investigation.

In conclusion, our novel mouse model has provided a powerful tool for probing the long-lasting effects of transient anti-NMDAR antibody exposure during a critical period of neurodevelopment, modeling the conditions seen in pediatric patients with NMDAR-AE. This experimental approach has allowed us to elucidate both cellular and network-level mechanisms that contribute to persistent sensory-motor deficits, even when autoantibody levels have declined. The schematic model presented in **Graphic Abstract** encapsulates these central findings and highlights the underlying mechanisms explored in this study.

This study emphasizes further the likely critical role of early diagnosis and intervention in pediatric NMDAR-AE. The implications of our work provide a compelling rationale for further studies in human patients to confirm these morphological changes and their correlation with the severity and persistence of behavioral deficits. The insights gained from this study lay a valuable foundation for future investigations into potential therapeutic interventions, ultimately advancing our understanding of NMDAR-AE and improving patient outcomes.

## Methods

### Sex as a Biological Variable

Both male and female mice were included across all experiments in this study to investigate potential sex-specific differences.

### Experimental Model and Subject Details

Floxed *Grin1* allele (Stock #005246), EMX1-Cre (Stock #005628) and Ai14 Cre reporter allele (Stock # 007914) were obtained from Jackson Laboratories (Bar Harbor, ME, USA). Floxed *Grin2a* and *Grin2b* alleles were provided by the laboratory of Prof. Roger Nicoll. Wild-type CD1 mice were obtained from Charles River Laboratories. Male and female embryos at embryonic (E) 15.5 were used for the *in utero* electroporation, and pups between postnatal day 0 (P0) to 14 (P14) for the experiments.

#### *In utero* Electroporation

DNA solution including the plasmid, and 0.04% fast green was injected into the medial region of the lateral ventricle of the embryonic brain with a glass micropipette. Electrical pulses then were delivered to embryos by electrodes connected to a square-pulse generator (ECM830, BTX). For each electroporation, five 35-V pulses of 50ms were applied at 1s intervals. After the electroporation, the uterus was returned to the abdominal cavity, followed by suturing of the abdominal wall and skin. Mice were perfused at different postnatal stages using 4% paraformaldehyde followed by post-fixed overnight and incubation in 30% sucrose at 4°C. 35 μm-thick coronal sections were obtained using cryostat sectioning.

#### Hippocampal Neuronal Cultures

Hippocampi from male and female *Emx1^cre/+^*; *Grin1^fl/fl^*; *Ai14^fl/fl^* and littermate *Emx1^cre/+^*; *Grin1^wt/wt^*; *Ai14^fl/fl^* mice were dissected at P0-P1 and incubated with trypsin at 37° C for 15 min. Cells were then dissociated by trituration with fire-polished Pasteur pipettes. Neurons were plated on poly-D-lysine (Sigma-Aldrich) and mouse laminin (Invitrogen) coated 12 mm glass coverslips (Warner Instruments) at 5x10^4^ cells per well in a 24-well plate in plating media (minimal essential medium [MEM], 10% FBS, 0.5% glucose, 1 mM sodium pyruvate, 25 µM glutamine, and 50 units penicillin/streptomycin). After 4 h, medium was changed to Neurobasal medium supplemented with B-27 (Gibco), GlutaMAX (Gibco), and 50 units pen/strep (Gibco). Cultures were maintained in an incubator with 5% CO2 at 37°C, and half of the medium was replaced as needed.

#### Hippocampal Slice Cultures Preparation and Treatment

Hippocampal slices were procured from rats aged between 6 to 8 days, following the protocol established by Stoppini et al.,(69). Each slice was subsequently prepared for patch-clamp recordings. Post-preparation, the slices were treated with 1 µl of the mAb3 monoclonal antibody solution, at a concentration of 2.0 µg/µl. This effectively delivered 2.0 µg of mAb3 to each slice. To allow for ample interaction between mAb3 and the NMDAR subunits, the slices were incubated with this antibody for a duration of 24 hours.

#### Intraventricular Injection

Antibodies were injected into the lateral ventricular of pups by a glass pipette with a sharp bevel at 45 degrees (BV-10 Micropipette Beveler, Sutter instrument). The diameter of the pipette tip was ∼40-80μm (70). The concentrations for antibody injections were 2.0μg/μl for mAb3 and Hum IgG. Antibodies were injected twice daily, and the injection volume was 0.8-1μl for each injection.

#### Methods to Prevent Bias

Mouse pups delivered from one dam were randomly assigned for injection. Pups from different dams at the same time were grouped for the experiment. Pups were raised by the mother until postnatal 28 days. After weaning, mice with different treatments were mixed-housed. The investigator was blinded to group allocation and data analysis.

#### Slice Preparation and Imaging

Mice were perfused with saline followed by 4% paraformaldehyde in phosphate buffered saline (PBS), pH 7.4. Brains were removed from mice and post-fixed in 4% paraformaldehyde overnight before being placed in 30% sucrose solution. The brains were then cut into 12-µm, 35-µm, and 200-µm sections with a cryostat (Leica VT1200S). Sections were imaged by Zeiss Axioscan Z.1 (Zeiss, Thornwood, NY, USA) with a 20X objective. Confocal images were taken by Zeiss LSM880 (Zeiss, Thornwood, NY, USA) with 20X objective, 63X oil objective, and 100X oil objective.

#### Tissue-Based Immunofluorescence

Mice were perfused with saline followed by 4% paraformaldehyde in phosphate buffered saline (PBS), pH 7.4. Brains were removed from mice and post-fixed in 4% paraformaldehyde overnight before being placed in 30% sucrose solution. The brains were then cut into 12-µm sections with a cryostat (Leica, VT1200S). Non-specific binding was blocked by adding 5% normal goat/donkey serum during pre-incubation and incubations in 1x PBS containing 0.05% TritonX-100. The primary antibodies were applied overnight at 4°C. Secondary antibodies were applied for 1-2 hours at 4 degrees and nuclei were stained with DAPI. Slides were mounted with Prolong Gold Anti-fade Mountant (Invitrogen, P36930).

#### Immunoprecipitation

Adult mice at P40 were used for immunoprecipitation. A single mouse brain was homogenized in homogenizing buffer (1% Triton X-100, 150 mM NaCl, 50 mM Tris-HCl pH 7.5, 2 complete protease inhibitor tablets (Roche), 1 PhosSTOP phosphatase inhibitor tablet (Roche)) with Dounce homogenizer on ice. The lysate was then centrifuged at 10,000g for 10 min at 4°C. For each IP, 300μl of supernatant was transferred into a labeled IP tube, and 700 μl of lysis buffer was added to achieve ∼2 μg/ μl brain lysate. 30μg monoclonal antibody (mAb) or ChromoPure IgG control (Sigma) was added to each IP tube and mixed by rotator for 4 hr at 4°C. Then, 50uL of equilibrated magnetic protein A/G beads (Thermo Scientific) were added to each IP reaction and mixed by rotator for 1 hr at 4°C. Afterward, beads were washed with cold lysis buffer three times. At last, 50 μl sample buffer was added to beads to elute the protein and the IP elution was then subjected to western blotting by SDS-PAGE of IP eluates on a 4-12% Bis-Tris gel.

### Quantification and Statistical Analysis

#### Callosal Axon Density Analysis

Sections were imaged by Zeiss Axioscan Z.1 (Zeiss, Thornwood, NY, USA) with 20X objective over whole brain section. Each image was made up by the compression of three slices in 4μm Z-stack. For each brain, only one section was chosen for data quantification. The callosal axon density (fluorescence density) in S1 was quantitatively analyzed by ImageJ software. See the details in Zhou et al., (31)

#### Individual Axon Terminal Tracing

200-μm coronal section of P120 mouse brain were imaged by Zeiss LSM880 (Zeiss, Thornwood, NY, USA) with 20X objective. 3D confocal image of S1 callosal axon terminals loaded in Imaris 9.8 (Oxford Instruments). The morphology of individual callosal axon terminals was tracked manually by combining Autopath function in Imaris. All calculations on traced callosal axon terminals were automatically done by Imaris.

#### Study Approval

All animal experiments were conducted in accordance with the regulations of the National Institutes of Health and were approved by the University of California San Francisco Institutional Animal Care and Use Committee (IACUC).

All human studies were approved by the University of California San Francisco Institutional Review Board (IRB). Informed consent was obtained from all participants prior to inclusion in the study.

#### Statistics

Statistical analyses were performed using Prism 9 (GraphPad Software, San Diego, CA) and R version 4.1.0 with packages such as ggplot2. Data are presented as scatterplots with mean[±[standard error (SE). The statistical tests used in this study included two-tailed student T-test, one-way and two-way analysis of variance (ANOVA), Mann-Whitney tests, Kruskal-Wallis tests, and Dunn’s multiple comparisons tests. Specific tests applied to each experiment are indicated in the respective figure legends. p <[0.05 was considered statistically significant.

## Data Availability

All underlying data supporting the findings of this study are provided in the "Supporting Data Values" Excel file accompanying this manuscript. This file includes all raw data for each figure presented in the study. Additional data and supporting analytic code are available from the corresponding author upon reasonable request.

## Author Contributions

J.Z. and S.J.P. designed the research. A.L.G., R.L., and M.R.W. developed the efficient single-cell isolation protocol, isolated the patient’s CSF B-cells, and produced the patient-derived monoclonal anti-NMDAR antibodies. C.M.B., R.L., T.T.N., B.T., and J.Z. verified these antibodies. C.M.B., R.L. and X.Z. performed the IP-MS and analyzed the data, including graph production. C.C., X.C., J.J.P., and R.A.N. conducted and analyzed the patch-clamp recording data. P.H., K.L., and J.J.P. assisted with MEA EEG surgery and recordings, while Y.L. and C.W. analyzed the MEA EEG data and graph production. M.L., D.N., and J.T.P. carried out the ex vivo network electrophysiology recordings, data analysis and graph production. J.Z. conducted most of the experiments, including mouse surgeries, behavior tests, anatomical studies, and single neuronal morphology analysis. J.Z., C.Z., and H.W. analyzed the behavioral and morphology data. The manuscript was written by J.Z., C.M.B., M.R.W., and S.J.P.

## Acknowledgements

We extend our gratitude to individuals and institutions that have contributed significantly to this work: **Technical Support and Discussion:** We extend our gratitude to Jeffrey Simms at the Gladstone Behavior Core for his invaluable assistance with experimental design and data analysis of behavioral tests. We also thank Dr. Dan Wang from Dr. Dena Dubal’s lab at UCSF and Dr. Wenjie Mao from Dr. Lennart Mucke’s lab at the Gladstone Institute for their support with behavioral tests and insightful discussions. Our appreciation goes to Blaise Ndjamen at the Gladstone Histology and Light Microscopy Core for his expertise in confocal imaging and Imaris data analysis, and to Dr. Mao Ye from UCSD for his guidance and discussions on NMDAR-mediated miniature excitatory postsynaptic currents.

## Data Analysis and Manuscript Preparation

Dr. Qin Ma from Dr. Jorge Oksenberg’s lab at UCSF, and Dr. Wenqing Li from UCSD provided invaluable manuscript editing support. We also thank our lab member Dr. Daniela Maria de Sousa Moura for her excellent assistance with data graphing.

## Fellowships and Awards

C.M.B. is grateful for the support from the Hanna H. Gray Fellowship, HHMI, and the University of California President’s Postdoctoral Fellowship Program. A.L.G. was supported by the NMSS Clinician-Scientist Development Award and the Kathleen C. Moore Postdoctoral Fellowship. M.R.W. was supported by Westridge Foundation.

## Patient Contribution

Special thanks to the patient whose participation was crucial for this research.

