## Supplemental information for "Disrupted callosal connectivity underlies long-lasting sensory-motor deficits in an NMDA receptor antibody encephalitis mouse model"

⸸ equal contributions; *, corresponding authors

**Supplementary information**

**Methods**

**Methods for Making Monoclonal Antibody from anti-NMDAR Encephalitis Patient**

*Patient Recruitment and Eligibility Criteria*

Inclusion criteria for this study was enrollment in “Immunological Studies in Multiple Sclerosis, Neuromyelitis Optica, Sarcoidosis, or Immune Mediated Central Nervous System Disorders” and diagnosis of antibody-positive NMDA receptor encephalitis. The patient is a 21-year-old Hispanic male who was experiencing his second attack of anti-NMDAR encephalitis. His first attack was treated with steroids and IVIg and he was then able to stop immunotherapies. Three years later, he became encephalopathic for a second time and was again found to have detectable anti-NMDAR antibodies in the CSF. The patient consented to donate CSF and blood, and patient demographics and clinical history were obtained as part of the above study. All studies were approved by the UCSF Institutional Review Board.

*Patient clinical history*

The research subject initially came to medical attention when he was 18 years-old with 6 weeks of behavioral changes, including becoming uncharacteristically short tempered, verbally abusive and paranoid. The symptoms progressed to more severe confusion, disorientation and the development of threatening auditory hallucinations which precipitated violent acts against himself and family members. He also developed dysarthria and diffuse tremors. His initial cerebrospinal fluid (CSF) exam was significant for 15 white blood cells/µL (80% lymphocytes, 20% monocytes) (normal 0-5 cells/µL), 0 red blood cells/µL (normal 0 cells/µL), protein 31 mg/dL (normal 15-45 mg/dL), glucose 53 mg/dL (normal 40-70 mg/dL) with multiple unique oligoclonal bands. EEG showed intermittent rhythmic delta activity maximally present in the left temporal and right occipitoparietal lobes. An extensive neoplastic work-up, including testicular ultrasound, whole body positron emission tomography-computed tomography (PET-CT), was unrevealing. The Mayo Clinic Laboratory CSF NMDA receptor antibody test was positive, and the subject received methylprednisolone, intravenous immunoglobulin and one cycle of rituximab.

The subject ultimately made a significant recovery and returned to work and living independently. He did not continue rituximab or other immunosuppressive therapy. Three years after his initial presentation, he was re-admitted to the hospital with worsening behavioral symptoms, including increased aggression and drug use, and worsening aphasia. His CSF was again inflammatory with 17 white blood cells/uL (78% lymphocytes, 12% neutrophils, and 10% monocytes), 32 red blood cells/uL, protein 25 mg/dL, glucose 68 mg/dL, multiple unique oligoclonal bands and an elevated IgG index of 0.9 (normal 0.3-0.6). His CSF NMDA receptor antibody was again positive. Surplus CSF from this time point was used for single cell B cell isolation and monoclonal antibody production. MRI brain with and without gadolinium did not show any parenchymal abnormalities, and repeat tumor screening was negative. The subject again improved clinically with a combination of intravenous immunoglobulin, methylprednisolone and rituximab.

*Patient Cerebrospinal Fluid Collection, Processing, and Fluorescence-activated cell sorting (FACS)*

12 ml of fresh CSF was centrifuged at 400g x 15 minutes at 4°C to separate a cell pellet from the supernatant. The supernatant was removed immediately to avoid cell dispersion, and care was taken to minimally move the source 15ml conical tube. Wide bore pipette tips were used when handling cells to preserve cell integrity.

Destination FACS tubes (1.5ml Eppendorf) were coated overnight with 100% FBS to minimize cell adhesion. Destination tubes were filled with 450μl of cold D-PBS + 15% FBS and placed lid-open in front of a de-static bar prior to placing them in the cell sorter. Cell samples were blocked with FcR Block (Miltenyi Biotec) and stained with fluorescent antibodies to cell surface markers: IgD Brilliant Violet 421 (Biolegend 11-26c.2a), CD20 FITC (Beckman Coulter B9E9), CD38 PerCPCy5.5 (BioLegend HIT2), CD3 PE-Cy7 (Beckman UCHT1), CD138 PE (Miltenyi 449), CD27 APC (eBioscience O323), CD19 APC-Alexa750 (Beckman J3-119). CSF B cell subsets were sorted on a Beckman MoFlo Astrios FACS sorter as previously described (1). CD27hi plasmablast/plasma cells (CD27hiPC: CD19+IgD-CD27hi), the vast majority of which are also CD38+, were then used for single-cell printing.

*Single Cell Printing of FACS-sorted CSF cells*

CD27^hi^PC cell suspensions were diluted to a 5% FBS concentration immediately prior to being centrifuged at 300g x 5min at 4**°**C. Pelleted cells were gently resuspended in 5-20μl of supernatant then placed on ice in a vacuum chamber for 5min. A ~5μl droplet of cells was pipetted onto a hydrophobic slide and aspirated for single cell printing by the CellenONE/sciDROP PICO system (Cellenion, Lyon, France and SCIENION, Berlin, Germany). Single cells were detected by the CellenONE and dispensed into wells of a 384-well PCR plate filled with 2μl/well of lysis buffer as described in the Smart-seq2 protocol (2).

*Single cell immune repertoire sequencing and monoclonal antibody production*

Cell-containing PCR plates were briefly vortexed and centrifuged at 2000 rpm x 1 minute, placed on dry ice and immediately stored at -80**°**C. We then performed RT-PCR using template switching and total cDNA amplification and bead clean as previously described (2). This was followed by targeted PCR of the heavy and light chains using primers to the leader sequences and constant region 1, followed by nested PCR as previously described (3). Products were sequenced via Sanger sequencing. Mapping of germline V, D, J and CDR3 regions was performed using IgBlast and IMGT V-Quest. Alignments were generated in Geneious (Biomatters). Recombinant monoclonal antibodies were produced with a high throughput method using a mammalian cell line (Tuna293) at LakePharma (San Carlos, CA), with mass spectrometry validation that the monoclonal antibody amino acid sequence matched the patient-derived Sanger sequence.

**Plasmid:** The ubiquitin-EGFP plasmid used was from a previous study (4).

**Antibodies: Commercial Antibody Table:**

| **Primary Antibody** | **Vendor** | **Assay** |
| --- | --- | --- |
| Goat anti-EphB2 | R&D, AF467 | TBIF 1:50 |
| Rat anti-GFAP | Invitrogen, 13-0300 | CBA 1:200 |
| Rabbit anti-HOMER1 | Synaptic System, 160 003 | CBA 1:1000 |
| Mouse anti-MAP2 | Millipore Sigma, MAB3418 | CBA 1:250 |
| Rabbit anti-GluN1 | Alomone, AGC-001 | CBA 1:300 |
| Rabbit anti-SYN1 (Synapsin) | Synaptic Systems, 101002 | CBA 1:1000 |
| Rabbit anti-β-Tubulin III (Tuj1) | Sigma-Aldrich, T2200 | CBA 1:2000 |
| Human IgG | Jackson ImmunoResearch,  009-000-003 | IP 30µg/700ul  Injection 1.6µg |
| **Secondary Antibody** | **Vendor** | **Assay** |
| Donkey Anti-Human IgG  (Alexa Fluor 488) | Jackson ImmunoResearch,  709-545-149 | TBIF 1:1000  CBA 1:1000 |
| Donkey Anti-Human IgG  (Alexa Fluor 594) | Jackson ImmunoResearch,  709-585-149 | CBA 1:1000 |
| Donkey Anti-Mouse IgG  (Alexa Fluor 488) | Jackson ImmunoResearch,  715-545-150 | CBA 1:1000 |
| Donkey Anti-Rabbit IgG  (Alexa Fluor 594) | Jackson ImmunoResearch,  711-585-152 | TBIF, CBA 1:1000 |
| Donkey Anti-Mouse IgG  (Alexa Fluor 594) | Jackson ImmunoResearch,  715-585-150 | CBA 1:1000 |
| Donkey Anti-Goat (Cy3) | Jackson ImmunoResearch,  705-165-003 | TBIF 1:1000 |
| Goat Anti-Human IgG (H+L)  (Alexa Fluor 488) | Jackson ImmunoResearch,  109-545-088 | TBIF 1:1000 |
| Goat Rabbit IgG (H+L)  (IRDye 800CW) | Li-Cor,  925-32211 | WB 1:20000 |

Patient-Derived Monoclonal IgG1Antibody Table

| Monoclonal Antibody | Assay |
| --- | --- |
| mAb1 | TBIF  CBA 10-18µg/mL  IP 30µg/700µl |
| mAb2 | TBIF  CBA 10-18µg/mL  IP 30µg/700µl |
| mAb3 | TBIF  CBA 10-18µg/mL  IP 30µg/700µl  IV Injection 2.0µg/µl |
| mAb4 | TBIF  CBA 10-18µg/mL  IP 30µg/700µl |

TBIF = Tissue-Based Immunofluorescence

IP = Immunoprecipitation

CBA = cell-based assay

WB = Western blot

IV = Intraventricular injection

**Live Neuron Staining**: At DIV 21, neurons were incubated with in-house monoclonal antibodies at a concentration of 18 µg/mL for 2 hours at 37°C. Following incubation, neurons were washed with warmed PBS and fixed with ice-cold 4% PFA in 0.1 M Na-acetate, pH 6.0 for 20 min at 4°C and washed thoroughly with PBS. Neuron-plated coverslips were blocked in 10% serum, 0.1% Triton-X-100 for 2 hours at room temperature. Primary antibodies were probed with donkey-anti-human Alexa Fluor 488 (Invitrogen) for 2 hours at room temperature and mounted in Prolong Gold (Invitrogen). Images were acquired using a Nikon Ti microscope equipped with an Andor Borealis CSU-W1 spinning disk confocal and Andor Zyla sCMOS camera (Nikon Core UCSF).

**Fixed Neuron Staining**

**Materials and Reagents:**

- Ice-cold 4% paraformaldehyde (PFA) in 0.1 M sodium acetate buffer (pH 6.0)
- Phosphate-buffered saline (PBS)
- 10% serum for blocking
- Monoclonal antibodies: mAb1 and mAb3 (10 μg/mL)
- Triton-X-100 (0.1%)
- Neuronal markers: MAP2, Tuj1, Synapsin 1, Homer 1, GFAP
- NR1 antibody (Alomone, AGC-001) conjugated to Alexa Fluor 647
- Prolong Gold Antifade (Invitrogen, P36930)
- Mix-n-Stain™ CF® Dye antibody labeling kit (Biotium, #92279)

**Protocol:**

1. **Fixation of Neurons:**

- At 21 days in vitro (DIV 21), cultured hippocampal neurons on coverslips were fixed with ice-cold 4% paraformaldehyde (PFA) in 0.1 M sodium acetate buffer (pH 6.0) for 15 minutes at 4°C.
- Following fixation, neurons were thoroughly washed with PBS to remove any residual fixative.

2. **Blocking and Primary Antibody Incubation:**

- To prevent non-specific binding while maintaining cell membrane integrity (non-permeabilized), coverslips were blocked with 10% serum for 30 minutes at room temperature.
- Fixed and blocked neurons were incubated overnight at 4°C with in-house monoclonal antibodies mAb1 and mAb3 at a concentration of 10 μg/mL. This ensured that the antibodies bound only to receptor sites accessible on the outer cell membrane without permeabilization.

3. **Secondary Antibody Incubation and Permeabilization:**

- After primary antibody incubation, coverslips were washed and then incubated with secondary antibodies for 1 hour at room temperature.
- Permeabilization was performed using 0.1% Triton-X-100 for 30 minutes at room temperature to allow intracellular staining.

4. **Co-staining with Neuronal Markers:**

- Neurons were stained with the following markers to confirm and compare the subcellular localization of mAbs:
  - **Dendritic Marker:** MAP2 (Millipore Sigma, MAB3418)
  - **Axonal Marker:** Tuj1 (Sigma-Aldrich, T2200)
  - **Presynaptic Marker:** Synapsin 1 (Synaptic Systems, 101002)
  - **Postsynaptic Marker:** Homer 1 (Synaptic System, 160 003)
  - **Astrocyte Marker:** GFAP (Invitrogen, 13-0300)
  - **NR1 Antibody:** (Alomone, AGC-001) conjugated to Alexa Fluor 647 using Mix-n-Stain™ CF® Dye antibody labeling kit (Biotium, #92279)

5. **Imaging:**

- Coverslips were mounted using Prolong Gold Antifade (Invitrogen, P36930).
- Imaging was performed with a Nikon Ti microscope equipped with an Andor Borealis CSU-W1 spinning disk confocal and an Andor Zyla sCMOS camera.

**Whole-Cell Voltage-Clamp Recordings in Acute Hippocampal Brain Slices**

Following the preparation of organotypic brain slice cultures from rats at P6-8 (5), slices were maintained, and 2 days later, patient-derived mAb was added and pre-incubated for 24 hours. Subsequently, whole-cell voltage-clamp recordings were obtained from pyramidal neurons in the CA1 region of the hippocampus, as per the method outlined by Schnell et al., 2002 (6). Neurons were identified based on their location and morphology. All recordings were conducted at a temperature of 20–25°C.

- **Internal Solution**: The internal pipette solution comprised 135 mM CsMeSO4, 8 mM NaCl, 10 mM HEPES, 5 mM QX314-Cl, 4 mM Mg-ATP, 0.3 mM Na-GTP, 0.3 mM EGTA, and 0.1 mM spermine. The osmolarity was adjusted to 290–295 mOsm, and the pH was buffered at 7.3–7.4.
- **External Solution**: The external solution contained 119 mM NaCl, 2.5 mM KCl, 4 mM CaCl2, 4 mM MgCl2, 1 mM NaH2PO4, 26.2 mM NaHCO3, and 11 mM glucose. This solution was continuously bubbled with a mixture of 95% O2 and 5% CO2 to maintain proper oxygenation and pH levels. To isolate excitatory postsynaptic currents (EPSCs), 100 μM of picrotoxin was added to the external solution to block inhibitory synaptic activity.
- **Recording of AMPAR and NMDAR EPSCs**:
- **Sequencing**: Sequential recordings were performed, starting with AMPAR EPSCs. Following this, NMDAR EPSCs were recorded sequentially.
- **AMPAR EPSCs**:
  - **Holding Potential**: –70 mV. This potential selectively measures AMPAR-mediated EPSCs, as NMDARs are inactive at this hyperpolarized voltage due to Mg2+ block.
  - **Events**: At least 50 EPSC events per cell were recorded to ensure reliable measurements.
- **NMDAR EPSCs**:
  - **Holding Potential**: +40 mV. NMDAR-mediated EPSCs were measured 100 ms after the stimulation artifact, a point at which the AMPAR component has fully decayed due to its faster kinetics. This ensures that the recorded currents are predominantly due to NMDAR activity.
  - **Events**: Similar to AMPAR EPSCs, at least 50 EPSC events per cell were recorded
- **Synaptic Stimulation and Current Recording**:
- Synaptic currents were evoked every 10 seconds using bipolar stimulating electrodes placed in the stratum radiatum.
- Current responses were collected with a Multiclamp 700B amplifier (Axon Instruments), filtered at 2 kHz, and digitized at 10 kHz. Cells with a series resistance larger than 20 MΩ were excluded from the analysis to ensure data quality.

**Calculation of AMPA/NMDA Ratios**

- Ratio Calculation: The AMPA/NMDA ratio was calculated by dividing the peak amplitude of AMPAR EPSCs (recorded at –70 mV) by the peak amplitude of NMDAR EPSCs (recorded at +40 mV) for each cell. This ratio provides a measure of the relative contribution of AMPAR and NMDAR components to the synaptic current.
- Data Analysis: The peak amplitudes for both AMPAR and NMDAR EPSCs were determined using Igor software. This software facilitated precise quantification and comparison of the EPSC amplitudes across different conditions.

**Method for Spontaneous NMDAR-mEPSC Recording**

1. **Hippocampal Neuron Culture Preparation**:

Hippocampal neurons were cultured from P0-1 *EMX1cre+/-; YFP* mouse pups. The neurons were specifically labeled with YFP fluorescence to identify excitatory neurons. Cultures were prepared according to established protocols (details in the Method section of **Hippocampal Neuronal Cultures**) and maintained under standard conditions until days in vitro (DIV) 14-21 before treatment and recording.

2. **mAb3 Treatment**:

Neurons were incubated with mAb3 at a concentration of 75 µg/mL for 24 hours prior to recording to assess the effects on NMDAR-mediated currents.

3. **Electrophysiological Recordings**:

- **Recording Conditions**: Cells were recorded at a holding potential of -80 mV using whole-cell patch-clamp recordings.
- **Intracellular Solution:** The patch pipette solution contained (in mM): CsMeSO4 135, NaCl 8, HEPES 10, Na-GTP 0.3, Mg-ATP 4, EGTA 0.3, QX-314 5, spermine 0.1, and 0.5 µM TTX. This solution was delivered directly into the patched cell.
- **Extracellular Solution**: Artificial cerebrospinal fluid (ACSF) containing the following (in mM): NaCl 119, KCl 2.5, NaH2PO4 1, NaHCO3 26.2, glucose 11, and CaCl2 2.5.
- **Recording Procedure**: Coverslips with neurons were transferred to the recording chamber pre-filled with the extracellular solution. No perfusion was used to avoid washout of mAb3 during the recording. YFP-expressing cells were identified under a fluorescence microscope, and whole-cell patch-clamp recordings were performed. NMDAR-mEPSCs were isolated after the addition of 0.5 µM TTX (to block action potentials), 15 µM NBQX (to block AMPA receptors), and 0.1 mM picrotoxin (to block GABA_A receptors) in the recording ACSF (7, 8).
- **Verification of NMDAR-mEPSCs**:
- **Mg²⁺ Block**: Spontaneous currents were only present at 0 Mg²⁺ and were blocked by 3 mM Mg²⁺, confirming that these currents were mediated by NMDAR.
- **APV Confirmation**: The addition of 50 µM APV at the end of each recording session resulted in the complete abolition of the currents, further verifying their NMDAR origin.

4. **Data Collection and Analysis**:

- **Data Acquisition:** Recordings were performed using a Multiclamp 700B amplifier (Molecular Devices), filtered at 1 kHz, and digitized at 10 kHz.
- **Event Detection and Analysis:** NMDAR-mEPSCs were identified based on their sensitivity to Mg²⁺ and APV. Events with a minimum threshold of 7 pA were selected for analysis.
- **Charge Transfer Analysis:** The area under the current deflection was calculated to quantify the charge transfer (Q transfer) of each mEPSC event (7). The averaged charge transfer over a 2-minute recording period per cell was used to quantify ionic movement during NMDAR-mediated mEPSCs.
- **Statistical Analysis:** Data were statistically analyzed using Prism software (GraphPad). NMDAR-mEPSC amplitude and charge transfer between control and mAb3-treated neurons were compared using unpaired t-tests, with p-values < 0.05 considered statistically significant.

**Mass Spectrometry (MS)**

IP samples were sent to University of California Davis Campus Mass Spectrometry Facilities for MS analysis.

**MS Data Analysis Process**

**Sample Preparation and Controls:** To ensure the specificity of mAb3, we performed targeted immunoprecipitation coupled with mass spectrometry (IP-MS). We prepared three biological replicates of mAb3-immunoprecipitated mouse brain lysate. Controls included empty beads to account for any non-specific protein binding, and mAbX, a monoclonal antibody targeting a different membrane protein (a zinc finger protein), used to control for background noise and specificity.

**Initial Filtering of Data:** Before normalization, we filtered the raw spectral counts from mAb3 immunoprecipitations to focus on proteins with substantial presence. Proteins with a mean raw spectral count of less than 10 across the mAb3 replicates were excluded. This step ensured that only proteins significantly represented in the sample were carried forward for further analysis.

**Normalization and Fold Change Calculation for Heatmap:** For the heatmap, spectral counts were normalized to bead-only controls to correct for non-specifically immunoprecipitated protein. The spectral count for each protein immunoprecipitated by mAb3 and mAbX was divided by the mean spectral count for bead controls to calculate their fold change relative to background. This fold change was log2 transformed and the most enriched proteins for mAb3 were displayed and the corresponding log2(FC)’s for mAbX were included as a comparator.

**Heatmap Generation:** A heatmap was generated using R to visually represent the log2 fold changes of proteins enriched in the mAb3 condition compared to the control mAbX. This graphical representation allows for easy identification of significant protein enrichments and depletions, showcasing the relative changes in protein abundance clearly and effectively.

**Normalization and Fold Change Calculation for Protein-Protein Interaction Network:** Separately, for the protein-protein interaction analysis, we averaged the spectral counts across three mAb3 replicates and controls, including mAbX and bead-only. Normalized weighted spectral counts were computed by calculating the fold change relative to bead control. That is, the mean spectral count for a given protein for mAb3 was divided by the mean spectral count of the bead controls. This fold change was log2 transformed.

**Protein-Protein Interaction Network Analysis**: We used STRING Protein analysis to identify potential interactions among the enriched proteins. The interaction data were then visualized using CYTOSCAPE, allowing us to construct a detailed protein-protein interaction network. This network highlighted the proteins that strongly interact with GRIN1 and GRIN2A, providing insights into the structural and functional relationships within the NMDAR complex.

**Supplementary Materials:** Detailed lists of proteins, including their spectral counts, fold changes are available in the tab labeled "**Supplementary Figure 2**" within the Excel sheet titled "**Raw Data for All Figures**." These details are also depicted visually in **Supplementary Figure 2**.

**Mouse Behavioral Tests**

All mouse behavioral tests were run in the Gladstone Behavioral Core. The use and care of the mice complied with the guidelines of the Institutional Animal Care and Use Committee of UCSF. All animals were maintained in 12 hr light/dark schedule and with access to food and water, ad libitum. Before behavioral testing, mice were transferred to the testing room and acclimated for at least 1 hour. All work surfaces were cleaned with Vimoba (100 ppm chlorine dioxide solution made from MB-10 Tablets, Quip Laboratories) before and after testing every day and with 70% alcohol between testing of different mice. The mice performed behavior were born on 8/16/2020. The following behavioral tasks were performed after they were 1 month old. The mice, born on 8/16/2020, were tested starting one-month post-birth, detailed as follows:

**Elevated Plus Maze Test (9/17/2020)**

An elevated plus maze is used to evaluate anxiety-related behavior. The maze consists of two open and two enclosed arms elevated 63 cm above the ground (Hamilton-Kinder, Poway, CA). Mice are placed at the intersection between the open and closed arms and allowed to explore for 10 min. Anxiety-related behavior was measured by the time and distance traveled in the open arm.

**Open Field Test (9/21/2020 - 9/28/2020)**

The open field test is used to measure locomotion and anxiety-related behavior. Mice are placed in the center area of the open field chamber (16 in x 16 in) with dim light in the dark phase for 10 min. Mice trajectory was recorded with photobeam assays (San Diego Instrument, San Diego, CA). The center area is an 8 in x eight in square in the center part of the chamber. Anxiety-related behavior was defined by the ratio of distance traveled in the center area compared to the total area. Total movements, rearing in the open field, and time spent in the center and periphery of the open field were recorded automatically for subsequent analysis.

**Nest Building Behavior Test (11/5/2020)**

Nest building is spontaneous home cage behavior of mice. For assessment of nest building behavior, group‐housed mice were transferred individually to new cages. A 5 × 5-cm white compressed cotton pad (Nestlets, Ancare) was placed in the center of the cage, and nest building behavior was assessed 2, 6, and 24 hours later. Composite nest building scores were assigned at each time point as follows: 0, nestlet untouched; 1, < 10% of nestlet shredded; 2, 10%–50% of nestlet shredded but nest was without shape; 3, 10%–50% of nestlet shredded and nest had shape; 4, 50%–90% of nestlet shredded but nest was without shape; 5, 50%–90% of nestlet shredded and nest had shape; 6, > 90% of nestlet shredded but nest was flat; and 7, > 90% of nestlet as shredded and nest had walls that were at least as tall as the mouse on > 50% of its sides. The task was performed on 11/5/2020.

**Burrowing Behavior Test (11/7/2020)**

Burring behavior is used to evaluate spontaneous home cage behavior of mice. A burrow is made of 200 mm long, 68 mm diameter tube. The open end of the tube is raised 30 mm by bolting two 50 mm machine screws through it, each 10 mm in from the end. The lower end of the tube is closed with a plug. Fill the burrow with 200 g food pellets for mice and place the burrow against the longer wall of a clean cage with a thin layer of bedding. The closed end of the burrow is against the back wall of the cage. Provide water but no extra food in the cage hopper for the mice. The test is started 3hr before the dark cycle. The displaced food pellets in the burrow were measured at 2hr and 24hr after cage-burrow setup.

**Rotarod Test (9/30/2020 - 10/2/2020)**

Rotarod is used to evaluate the motor coordination of mice. Rotarod was performed with a Med Associates ENV-575(M) apparatus, using a 3-day protocol. Day 1 involved three training runs at a fixed 16 rpm. Days 2 and 3 each involved 6 data runs, 3 in the morning and 3 in the afternoon. The data runs on days 2 and 3 used steady acceleration of the rod from 4-40 rpm over 5 min. The time was stopped at 5 min or when the mouse fell. Photobeams were interrupted by the experimenter if the mouse held onto the rod without walking for three full rotations. Each mouse was given three trials at least with a 15-min inter-trial interval and a maximum of 300 s per trial. The average latency to fall off the Rotarod was calculated.

**Balance Beam Test (10/5/2020 - 10/6/2020)**

The balance beam is used to evaluate motor coordination and balance. A 15” long plastic beam is suspended about 20” above the table, with an open platform at one end and a dark box with 1.5” square opening at the other end. The balance beam test was taken place over 3 days with 3 trials per day. The first day included two additional training trials. On the first training trial of the first day, mice were placed on the middle of the largest diameter beam facing a dark box and guided to the enclosure. On the second training trial on the first day, mice were placed on the end of the beam facing the dark box and guided to the enclosure. On the third to fifth trials (testing trials) on the first day, mice were placed on the end of the beam facing the dark box and traversed the beam unguided. On the second day, mice were placed at the end of the medium diameter beam, facing the dark box. On the third day, mice were placed at the end of the smallest diameter beam, facing the dark box. Latency to traverse the beam into the dark box, number of foot slips, and the number of falls were recorded. Trials were limited to a maximum latency of 60s.

**Pole Test (10/8/2020 - 10/9/2020)**

The pole test evaluates the ability of the mouse to grasp and maneuver on a pole in order to descend to its home cage. The vertical pole is 50 cm long (1 cm diameter) with a rough surface. The base of the pole was placed in the home cage. The pole test was taken place over 2 days. The first day included three facing down training trials and three facing up testing trials. The second day included three facing up testing trials. In the training trials, mice were placed facing down atop of the pole and the time to descend the pole was measured. In the testing trials, mice were placed upward atop of the pole. The latency for mice to turn around and climb down was recorded. Any slips or falls in both training and testing trials were also recorded. If a mouse falls at the beginning of the trial due to placement issues, then the mouse is replaced, and the trial is restarted.

**Grip Strength Test** **(11/17/2020)**

The Grip Strength test is a complement to behavioral tests for motor coordination and motor function. The Grip-Strength Meter (Ugo Basile) was used to measure the grip strength on the forelimbs and all limbs. Mice were placed over a base plate, in front of a grasping bar (either trapeze-shaped or the grid), whose height is adjustable. The grasping tool is fitted to a force sensor connected to the Peak Amplifier, which guarantees a reliable and automated detection of the animal response. The force transducer has a maximum applicable force of 1500g, with a resolution 0.1g.

**Inverted Grid Hang Test (11/19/2020 - 11/22/2020)**

The inverted grid hang test is used to measure the ability of mice to exhibit sustained limb tension to oppose their gravitational force. The apparatus consists of a 15x25 cm grid of fine wire mesh suspended 34 cm above the table on supports. The walls and rear of the supports are covered with smooth plastic boards to prevent escape, and a larger plastic board is placed atop the wire mesh to prevent a mouse from climbing around the edge. The tabletop was covered with layers of soft towels to prevent injury from a fall. Mice were placed on the wire mesh, gently shaken to ensure a grip, and the mesh then inverted onto the supports. Latency to fall was recorded within 180 s. Measurements were repeated three times.

**EEG Surgery and Recording System Utilizing a 30-Contact MEA Probe during Pole Test in Mice**

Our research involved the use of a 30-contact multielectrode array (MEA) probe from NeuroNexus (Mouse EEG_v2-H32) to investigate electroencephalographic (EEG) activity in mice while performing a facing down and facing up pole test.

Initially, mice were anesthetized using isoflurane (at 4% for induction and maintained at 1-2%) in an induction chamber, and then positioned in a stereotaxic frame. We continuously monitored the depth of anesthesia to ensure the animals' wellbeing. A heating pad was utilized throughout the surgery to maintain the animals' body temperature.

After induction of anesthesia, the fur over the surgical site was removed, and the scalp was sterilized using povidone-iodine solution and alcohol. A midline incision was made to expose the skull, which was cleaned meticulously to create a smooth, dry surface. Following the technique outlined in (9), three screws were placed on the surface as shown in **Supplementary Figure 12A**, and the Bregma was marked for aligning the MEA probe. A wire soldered to the screw on the left cerebellum was used to connect the reference electrode with the MEA probe. Utilizing a stereotaxic manipulator, the MEA probe was meticulously positioned on the surface of the mouse brain as detailed in **Supplementary Figure 12B**, and the remaining steps were completed as per the reference (9) to finish the MEA probe implantation. This method not only ensured the stability of the MEA probe for consistent and reliable EEG recording but also allowed for the probe to be reused in subsequent experiments. The reusability of the probe contributes to the cost-effectiveness and sustainability of the research protocol.

Upon completion of the surgery, the mice were allowed to recover with careful monitoring. Post-operative analgesics were given as necessary to relieve any potential discomfort.

We recorded the EEG signals while the mice were performing the facing down and facing up pole tests, using the Plexon system. The MEA probe was connected to a preamplifier, which was in turn linked to the main amplifier and data acquisition system. The Plexon system was calibrated according to the manufacturer’s guidelines to ensure accurate signal detection.

The EEG signals were recorded at a sample rate of 1000Hz for the duration of the pole tests. The recorded EEG signals are the clear signals with subtraction of the reference signal. This method effectively isolates the brain activity of interest by removing common noise, including muscle artifacts, which might otherwise influence the EEG readings. The acquired data was then stored for further analysis.

According to the published mouse brain map (10), we annotated different cortical regions with electrode numbers in different systems as shown in **Supplementary Figures 12C-12E**. For our analyses, we specifically focused on the somatosensory and motor cortical regions. We used three channels to represent each region to ensure robust data capture:

- Left Primary Somatosensory Cortex (S1): Channels 17, 19, and 27
- Right Primary Somatosensory Cortex (S1): Channels 6, 4, and 26
- Right Primary Motor Cortex (M1): Channels 8, 7, and 5

These channels are averaged for each respective brain region before performing cross-correlation coefficient analyses to assess functional connectivity.

Throughout the experimental period, we regularly monitored the animals to ensure adherence to ethical guidelines and to minimize potential distress.

**EEG Functional Connectivity Analysis**

All analyses were done using MATLAB (MathWorks, Natick, MA) as previously described (11, 12). The raw EEG recording data were converted into .mat files in microvolt units and 60 Hz notch filter was applied to remove powerline interference. EEG data were then clustered into canonical frequency bands (theta 4-10 Hz, beta 10-30 Hz, slow gamma 30-55 Hz, and fast gamma 55-100 Hz) (13) using a discrete-form finite impulse response bandpass filter. The attenuation in the stop band was set as 80 dB and the amount of ripple allowed in the pass band was 1 dB. The frequency differences between the start of the 1^st^ stop band and the start of the 1^st^ pass band and between the start of the 2^nd^ pass band and the start of the 2^nd^ stop band were both 0.5 Hz.

To determine the functional connectivity (neural coupling) between two hemispheres or brain regions, cross-correlation analysis was performed in MATLAB (xcorr function). The instantaneous amplitudes of the recorded EEGs from the left and right S1 or right M1 and S1 were considered as two discrete-time sequences *x(n)* and *y(n)* respectively. The cross-correlation coefficient $\hat{R}_{xy,coeff}$ was calculated by the xcorr function in MATLAB as the following formula (the asterisk denotes complex conjugation):

$$\hat{R}_{xy}\text{(m)}\text{=}\left\{ \begin{aligned} \sum_{n=0}^{N-m-1} x_{n+m}y_{n}^{*}, m\geq0, \\ {\hat{R}^{*}}_{yx}\text{(-m), m<0.} \end{aligned} \right.$$

$$\hat{R}_{xy,coeff}\text{(m)}\text{=}\frac{1}{\sqrt{\hat{R}_{xx}\text{(0)}\hat{R}_{yy}\text{(0)}}}\hat{R}_{xy}\text{(m)}$$

The averaged peak cross-correlation coefficient was retrieved by averaging the peak cross-correlation coefficient $\hat{R}_{xy,coeff-peak}$ across all the animals in the control group or treatment group during the particular trials.

**Slice Preparation for Neocortical Network Recording**

Coronal neocortical 400 μm – thick brain slices containing the S1 cortex were obtained using the Leica VT1200 microtome (Leica Microsystems) as described in previous studies (14-16). Slices were placed in an interface chamber at 34 °C and perfused at a rate of 2 mL.min-1 with oxygenated artificial cerebro-spinal fluid (aCSF) containing 126 mM NaCl, 2.5 mM KCl, 1.25 mM NaH2PO4, 1 mM MgSO4, 2 mM CaCl2, 26 mM NaHCO3, 10 mM glucose, and 0.3 mM glutamine equilibrated with 95 % O2 and 5 % CO2, pH 7.4.

**Data Acquisition**

Extracellular multi-unit recordings were obtained with a linear 16-channel multi-electrode array with 100 μm spacing between the electrodes (Neuronexus, A16x1-2mm-100-703) (14) placed in the S1 cortex as shown in **Figure 8A**. The orientation of the probe was such that electrode 1 was placed in layer 1 of the cortex. Position of recording array was visually checked for each recorded slice to confirm the location of the electrodes in the different layers. The anatomical location of the electrodes was typically as follows: electrode 1 in layer 1; electrodes 2, 3, 4 in layers 2/3; electrodes 5, 6, 7 in layer 4, and electrodes 8, 9, 10 in layer 5. The signals were sampled at 24.414 kHz. Signals were amplified at 10,000× and band-pass filtered between 100 Hz and 6 kHz using the RZ5 signal processor from Tucker-Davis Technologies (TDT, SCR_006495). Electrical stimuli (10, 20, 50, 100, 150, 200, 300, 400, 500, and 600 μA) were delivered to the corpus callosum with a concentric bipolar tungsten electrode (50–100 kΩ, FHC Inc.). Stimulation pulses (100 μs duration) were repeated nine times in a single recording spaced by 8 seconds.

**Quantification of Spikes**

Spikes were extracted using a custom Spike2 (SCR_000903) script that enumerated all negative deflections that fell through a threshold of -22.7 μV. The threshold was chosen because it reliably captured the spikes without false negatives. Spikes were analyzed over a defined interval from the time of the stimulus to 25 ms after, as indicated by orange horizontal bars in **Figures 8B, 8C**. The number of spikes was quantified for each recording channel (**Figure 8D**); then averaged between channels 2–4 to obtain average spike rates for layers 2/3, between channels 5–7 to obtain average spike rates for layer 4, and between channels 8–10 to obtain average spike rates for layer 5.

**Statistical Analysis**

All numerical values are given as means and error bars are standard error of the mean (SEM) unless stated otherwise. Data analysis was performed with SigmaPlot (SCR_003210), MATLAB (SCR_001622), Origin 9.0 (Microcal Software, SCR_002815), and GraphPad Prism 7 (SCR_002798)

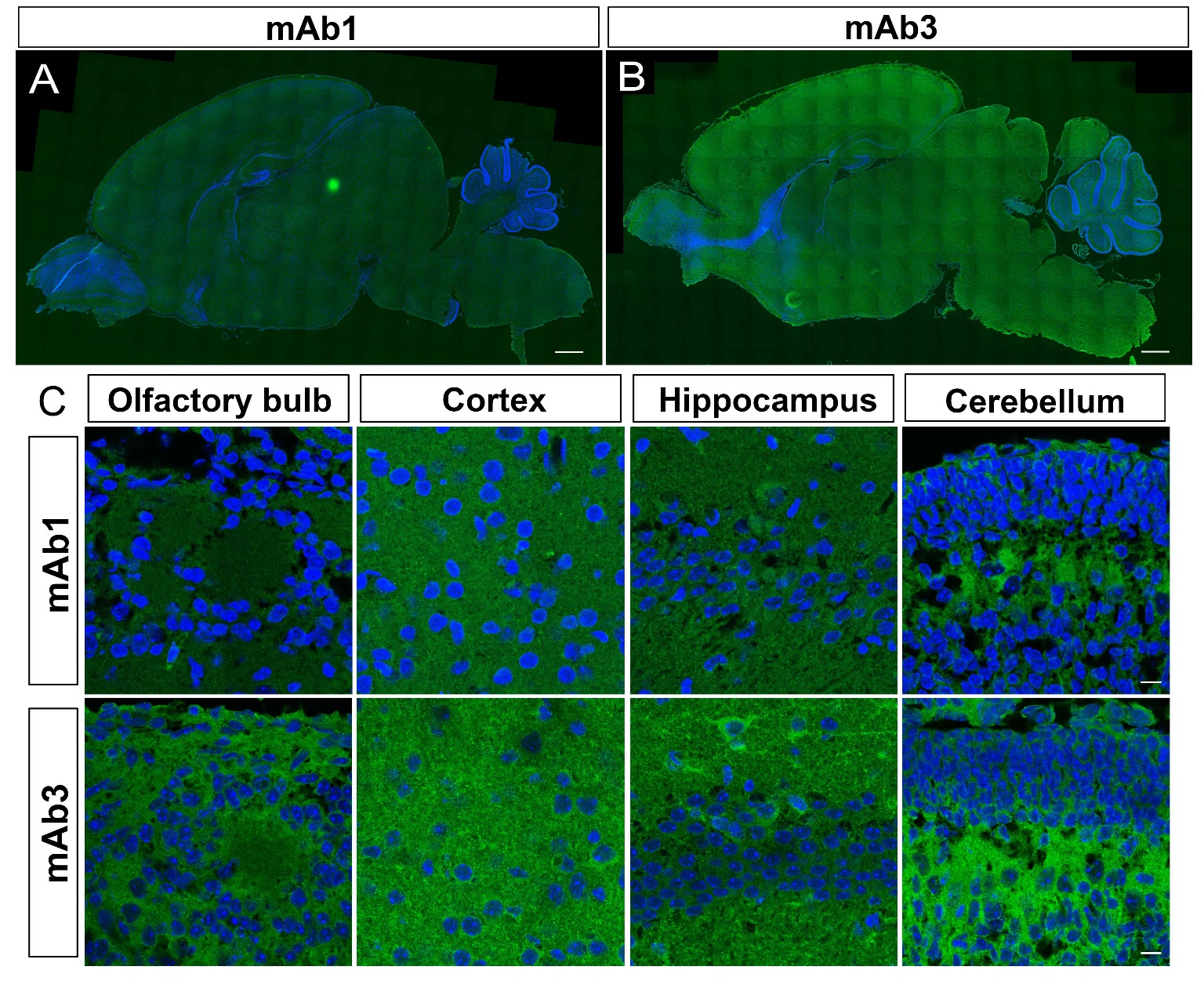

**Supplementary Figure 1: Immunostaining pattern of mAb1 and mAb3 at early developmental stage**. The immunostaining pattern mAb1 (A) and mAb3 (B) on sagittal sections of P8 mouse brains. (C) Immunostaining pattern of mAb1 and mAb3 in different brain regions. Scale bar: 500μm for images A, and B; 10μm for images in C.

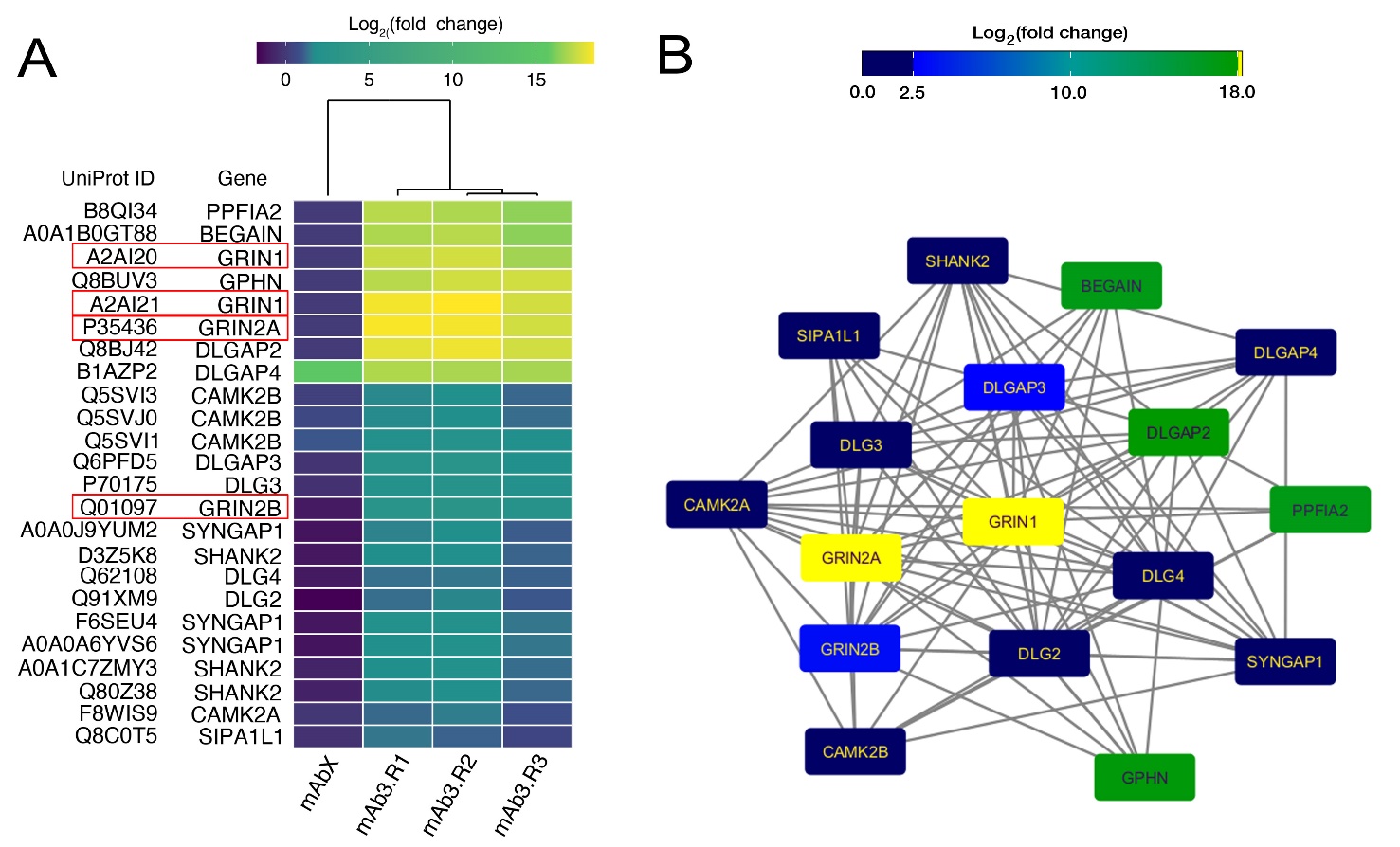

**Supplementary Figure 2**: **Enrichment of NMDAR complex proteins GRIN1 and GRIN2A in mouse brain lysate immunoprecipitated by mAb3.** (A) The heatmap displays the bead-normalized log2 fold change of spectral counts for mouse brain proteins immunoprecipitated by mAb3 (N = 3 technical replicates) and mAbX, which was raised against a zinc finger protein that localized on the cell membrane. Members of the NMDAR complex, notably GRIN1 and GRIN2A, are outlined in red. (B) Protein-protein interaction network for GRIN1 (log2FC = 18.79), GRIN2A (log2FC = 18.12), and other immunoprecipitated proteins. Their enrichment relative to bead controls is indicated by the color scale.

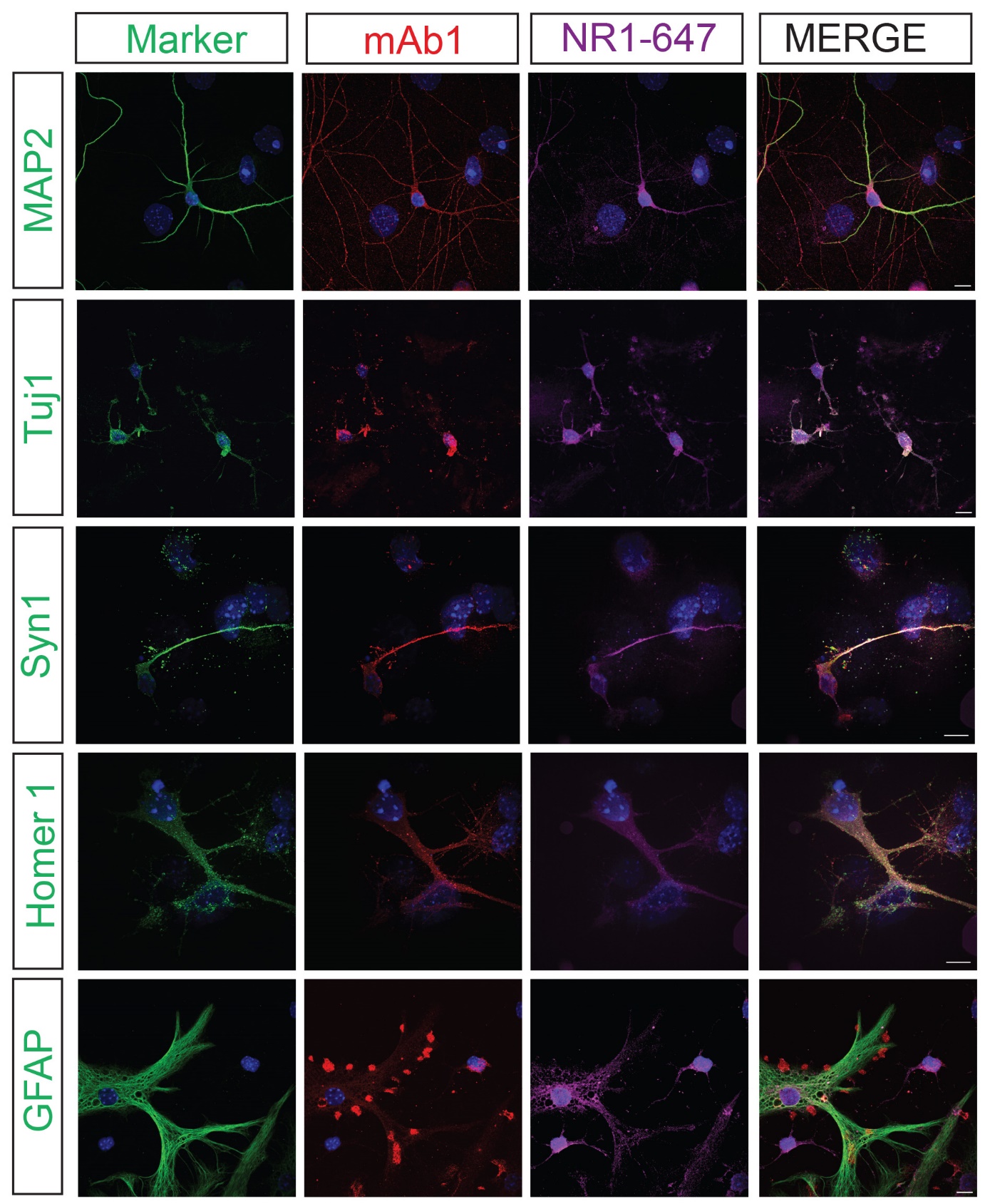

**Supplementary Figure 3: Subcellular Localization and Specificity of mAb1.** This figure demonstrates the specific subcellular localization of mAb1, which was initially applied to fixed, non-permeabilized cultured hippocampal neurons to ensure precise surface labeling. After the application of secondary antibodies to mAb1, cells were permeabilized, facilitating the introduction of neuronal and astrocytic markers, as detailed in the Methods section. The staining protocol included the dendritic marker MAP2, axonal marker Tuj1, presynaptic marker Synapsin 1 (Syn1), postsynaptic marker Homer 1, and astrocytic marker GFAP, in addition to a commercially conjugated anti-GluN1-647 (NR1-647) antibody. The organization of the figure is by rows, each representing a different marker (MAP2, Tuj1, Syn1, Homer 1, GFAP), and by columns displaying from left to right: the specific marker alone, mAb1 staining, NR1-647 antibody, and the merged images. mAb1 exhibits punctate staining patterns that significantly co-localize with MAP2, NR1-647, Syn1, and Homer 1, indicating its association with dendritic and synaptic sites. This co-localization with NR1-647 at these synaptic markers suggests mAb1’s involvement with NMDARs in synaptic regions. However, its interactions with the axonal marker Tuj1 are less pronounced, featuring discontinuous puncta that contrast sharply with the continuous staining of NR1-647, which only partially overlaps with mAb1. These observations are consistent with mAb1's punctate staining around nuclei in mouse brain tissue, without defining neuronal morphology. Intriguingly, mAb1 does not overlap with GFAP-positive astrocytes or co-localization with NR1-647 in astrocytes, it targets distinct clusters around the GFAP-positive astrocytes that do not overlap with NR1-647. This distinct expression pattern of mAb1, alongside neuronal and astrocytic markers, and contrasting with the anti-GluN1 subunit antibody NR1-647, further suggests that mAb1 interacts with NMDARs at synapses but likely recognizes a different NMDAR subunit than GluN1. Scale bar: 10 µm.

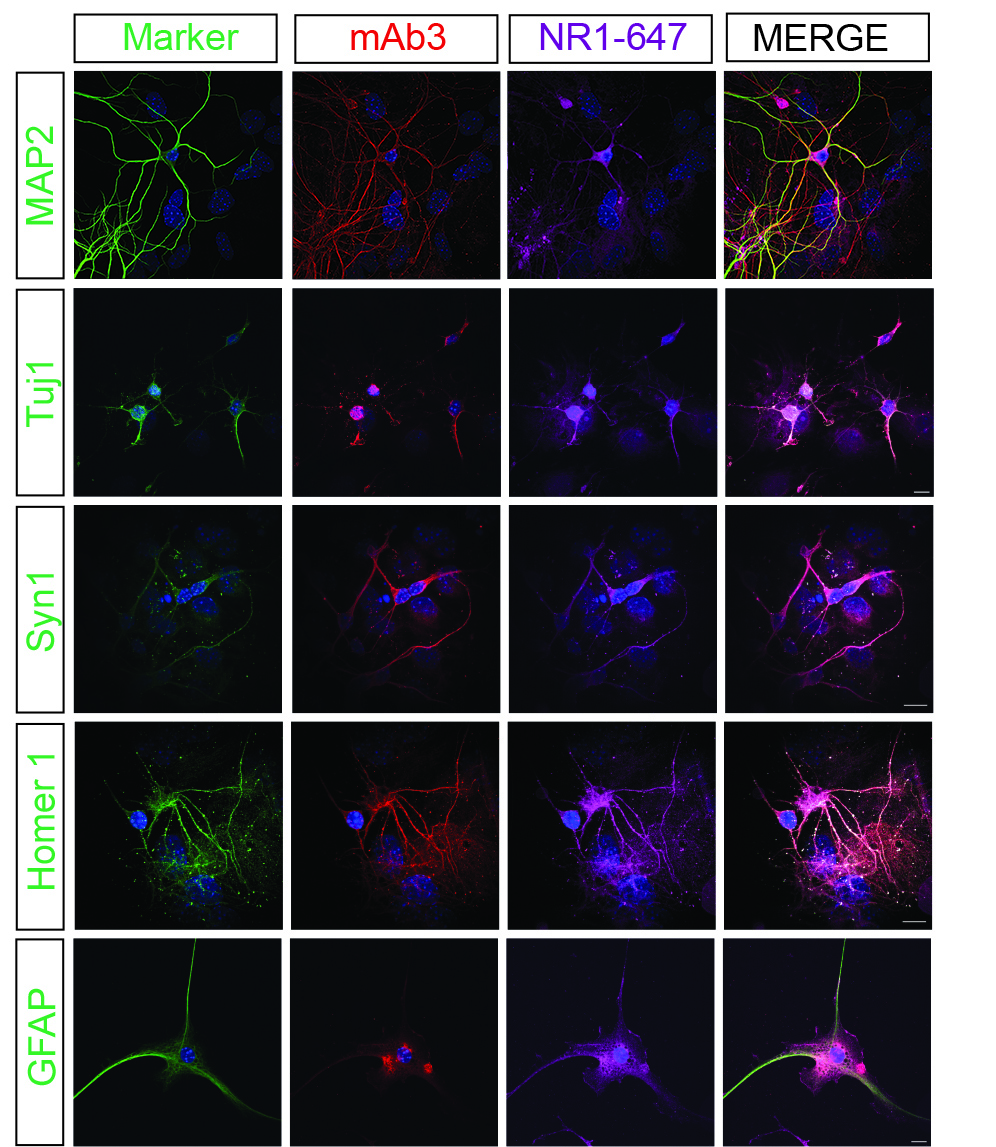

**Supplementary Figure 4: Subcellular Localization and Specificity of mAb3.** Following the same staining protocol as described for mAb1, this figure presents high-resolution images demonstrating the surface co-staining of mAb3 with a series of neuronal and astrocytic markers: MAP2 (dendritic), Tuj1 (axonal), Synapsin 1 (Syn1, presynaptic), Homer 1 (postsynaptic), and GFAP (astrocytic). The organization of the figure is consistent, with rows showing the progression from the specific marker alone, to mAb3 staining, to co-staining with the anti-GluN1-647 (NR1-647) commercial antibody, and finally the merged images. mAb3 exhibits a continuous distribution closely aligning with dendritic, axonal, presynaptic, and postsynaptic markers, consistent its expression in neuronal cell bodies and processes as observed in mouse brain. Extensive co-localization with the NR1-647 commercial antibody across all these neuronal markers underscores mAb3’s specificity for the GluN1 subunit of NMDARs. Notably, mAb3 also forms bright aggregates near GFAP-positive astrocyte cell bodies and processes that co-localize with the aggregates targeted by the NR1-647 commercial antibody, further reinforcing its specificity for the GluN1 subunit of NMDARs. Scale bar: 10 µm.

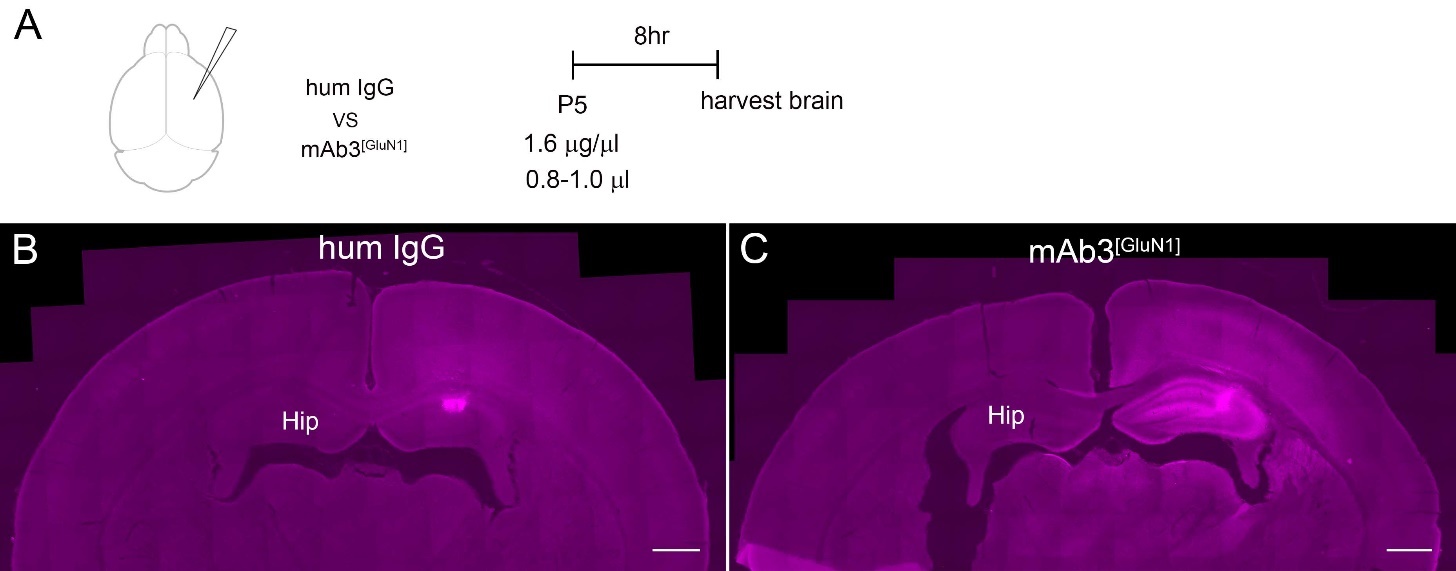

**Supplementary Figure 5: The diffusion territory of mAb3^[GluN1]^** **in mouse cortex after 8 hours of intraventricular injection.** (A) We injected mAb3^[GluN1]^ into the lateral ventricle at P5 and perfused the mouse 8 hours later. The Human IgG was injected into the same littermates as controls. We stained the mouse brain with anti-Human secondary antibody coupled to Cy5, which thus indicated where the antibodies had diffused in the mouse brain. (B) In control mice, the Human IgG had diffused to motor and sensory cortexes, but the signal was getting weaker. (C) However, the mAb3^[GluN1]^ injected brain still had strong signals in the cortex and hippocampus on the injection side. Scale bar: 500μm for all images. Hip: hippocampus.

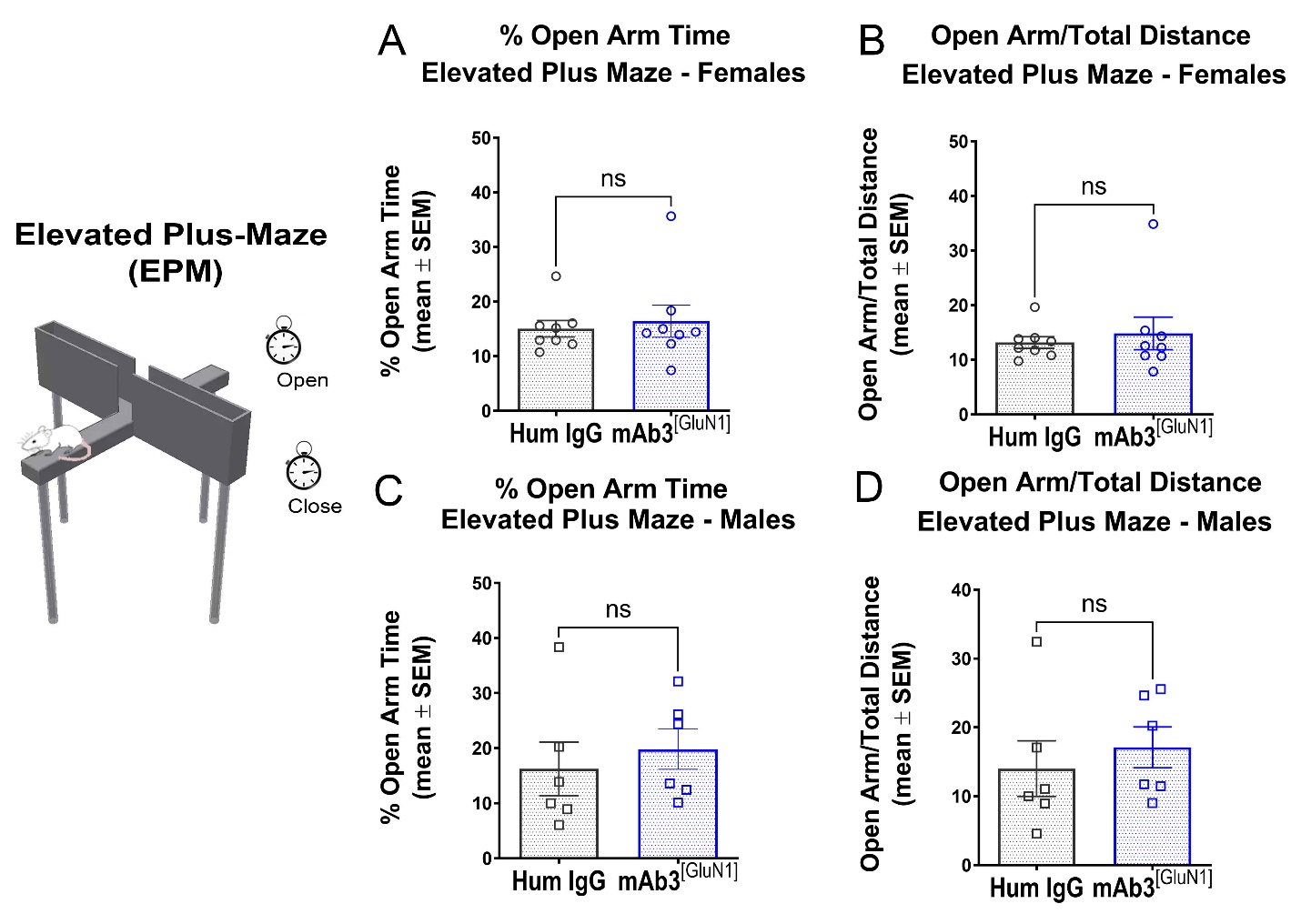

**Supplementary Figure 6: No increased anxiety in mAb3^[GluN1]^-treated mice.** (A, B) Time traveled in the open arm of Elevated Plus-Maze (EPM). There was no difference in time spent on the open arms between Human IgG and mAb3^[GluN1]^ treated female (A, P = 0.96) and male (B, P > 0.999) mice. (C, D) Distance traveled in the open arm of EPM. There was no difference in distance traveled on the open arms between Human IgG and mAb3^[GluN1]^ treated female (C, P = 0.39) and male (D, P = 0.31) mice. n = 6 to 8 per group. The p value represents the statistical difference between the two groups by using Mann-Whitney test.

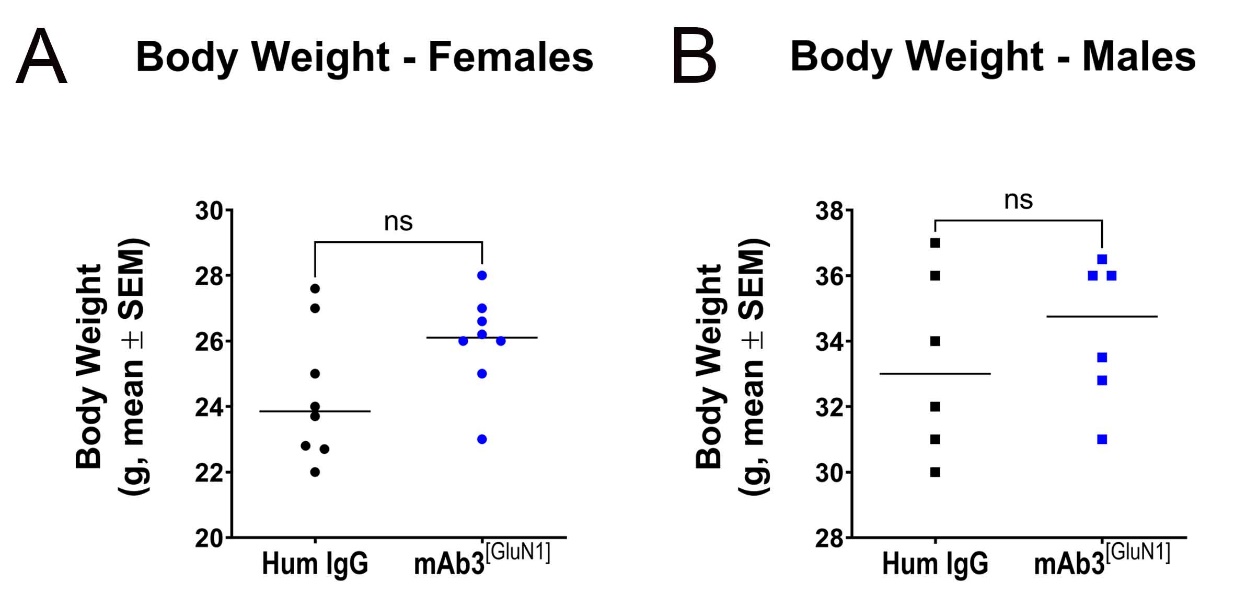

**Supplementary Figure 7: No body weight difference between groups**. Comparison of body weight in female (A, P = 0.12) and male mice (B, P = 0.62). Above statistics were based on Mann-Whitney test.

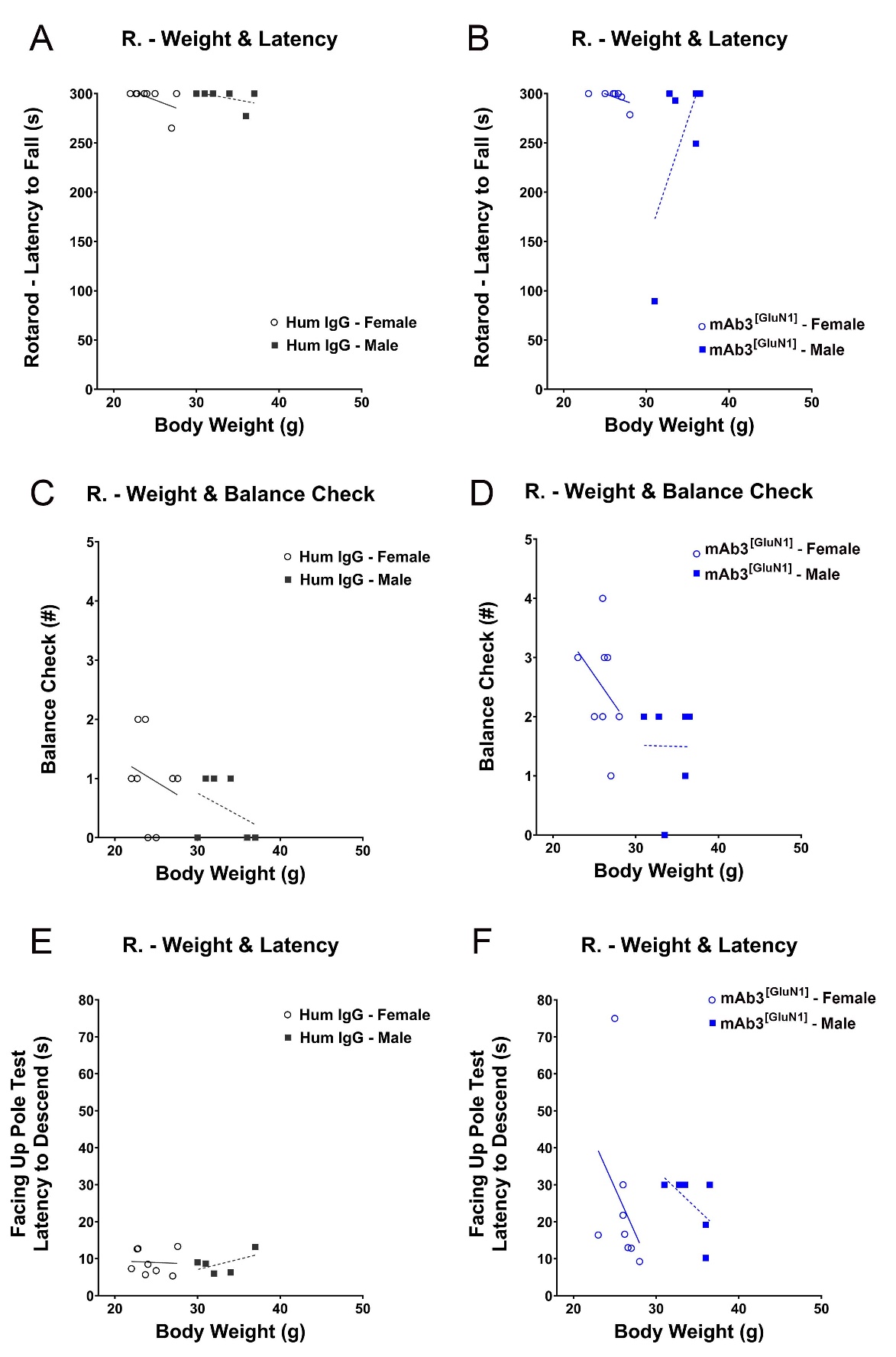

**Supplementary Figure 8: No correlation between body weight and motor performance.** (A, B) Correlation between weight and latency to fall (session 2 of rotarod). (A) Human IgG (Female: R^2^ = 0.27, P = 0.18 by linear regression; Male: R^2^ = 0.22, P = 0.35). (B) mAb3^[GluN1]^ (Female: R^2^ = 0.36, P = 0.11; Male: R^2^ = 0.44, P = 0.15). (C, D) Correlation between weight and balance check (trial 4 of balance beam). (C) Human IgG (Female: R^2^ = 0.05, P = 0.58; Male: R^2^ = 0.15, P = 0.44). (D) mAb3^[GluN1]^ (Female: R^2^ = 0.10, P = 0.44; Male: R^2^ = 0.0001, P = 0.98). (E, F) Correlation between weight and latency to descend (trial 2 of facing up pole). (E) Human IgG (Female: R^2^ = 0.003, P = 0.90; Male: R^2^ = 0.29, P = 0.35). (F) mAb3^[GluN1]^ (Female: R^2^ = 0.12, P = 0.40; Male: R^2^ = 0.31, P = 0.25).

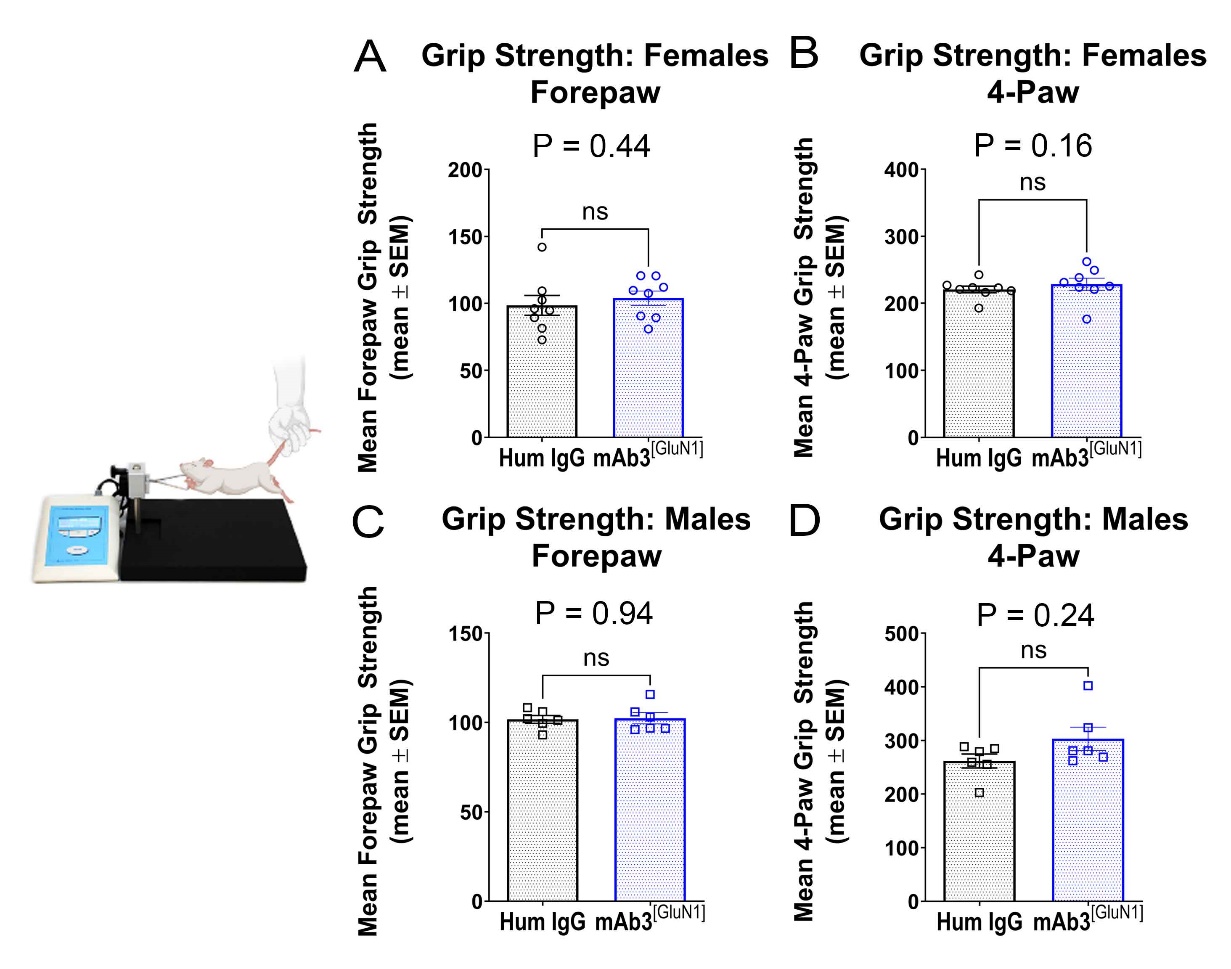

**Supplementary Figure 9: No impaired muscle strength in mAb3^[GluN1]^-treated mice.** (A, B) Grip strength in female mice. Human IgG served as controls. There was no difference in mean forepaw (A) or mean 4-paw (B) grip strength in female mice. (C, D) Grip strength in male mice. There was no difference of mean forepaw (C) or mean 4-paw (D) grip strength in male mice. The above statistics were based on Mann-Whitney test.

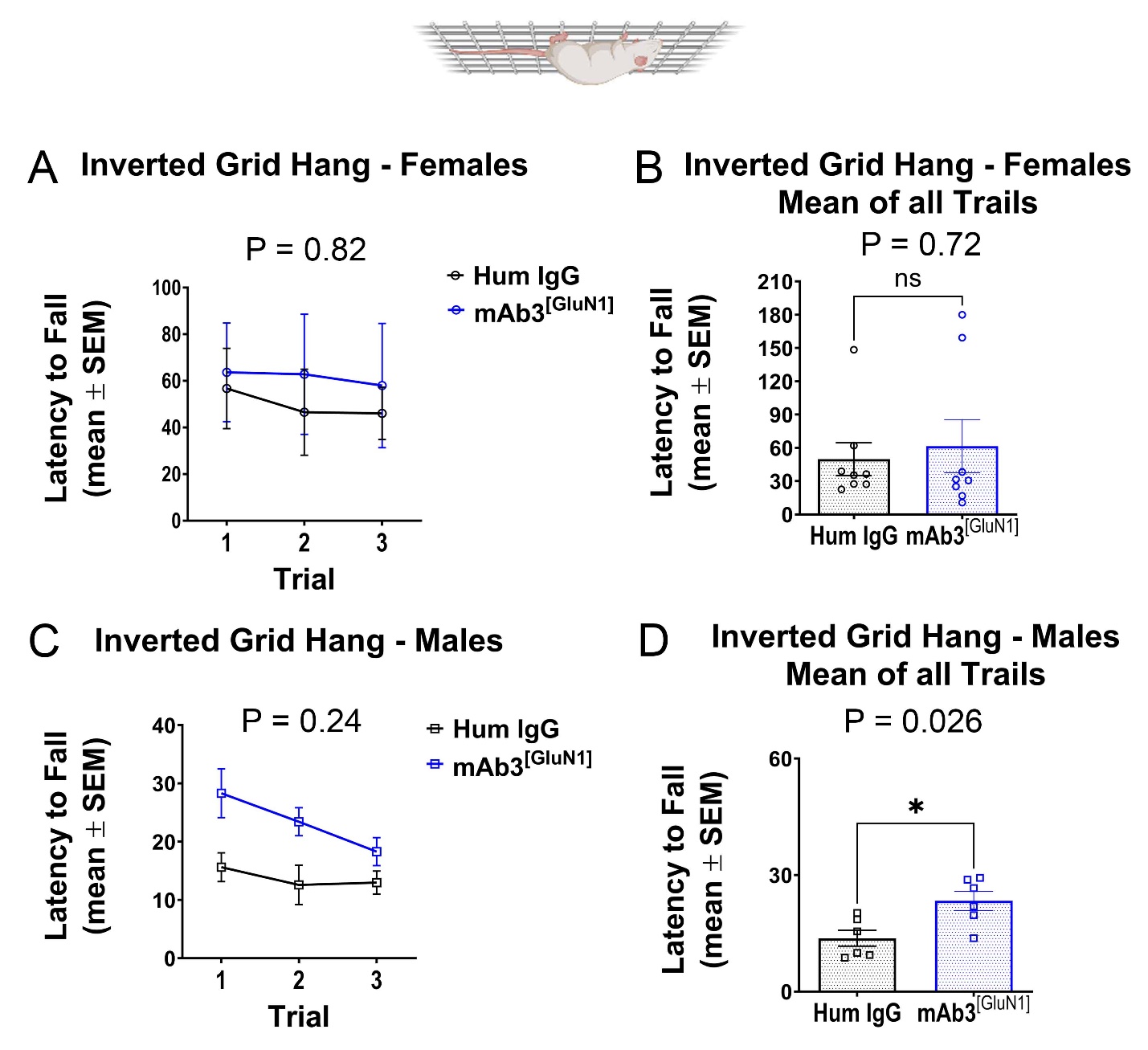

**Supplementary Figure 10: No impaired muscle strength and coordination in mAb3^[GluN1]^-treated mice.** (A, B) Inverted grid hang of female mice. Human IgG served as controls. There were no differences of latency to fall in female mice. (C, D) Inverted grid hang of male mice. mAb3^[GluN1]^-treated male mice took significantly longer to fall (B) compared to the control group, suggesting they exhibited better muscle coordination than controls. The above statistics were based on Mann-Whitney test.

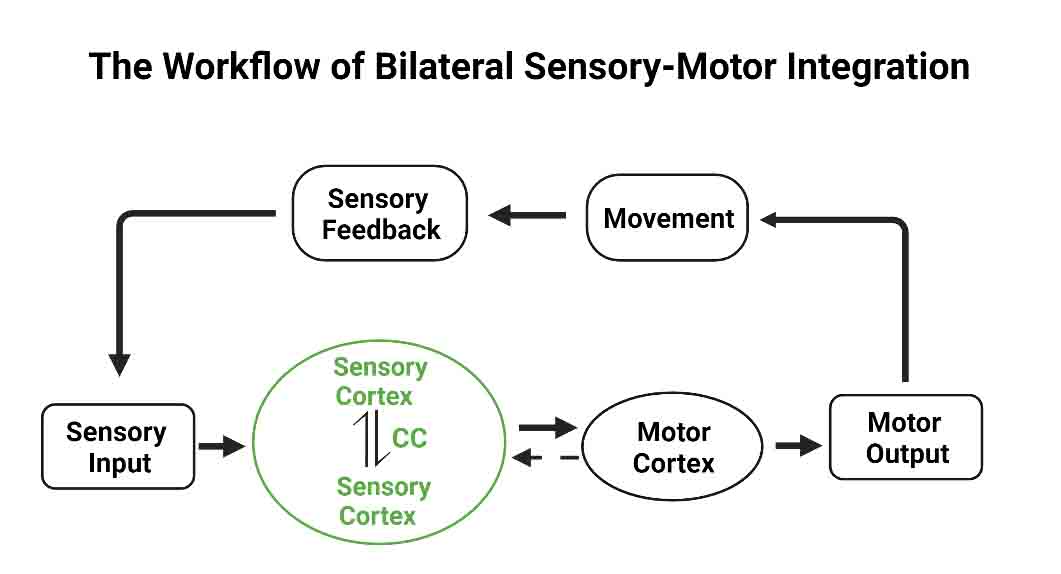

**Supplementary Figure 11: Model of sensory-motor integration**. Schematic illustration of the sensory-motor integration process. The diagram begins with sensory input, which is relayed to the primary somatosensory cortex (S1) for interpretation. Inter-hemispheric connections via callosal axons enable integration and coordination of sensory information across both hemispheres, which is crucial for bilateral sensory processing. Intra-hemispheric connections between S1 and the primary motor cortex (M1) facilitate the translation of sensory information into motor commands within the same hemisphere, essential for executing movements. Motor responses planned in M1 lead to movement, which generates sensory feedback sent back to S1, forming a feedback loop for continual adaptation of motor output based on sensory input. This diagram highlights the critical roles of both inter-hemispheric (S1-S1) and intra-hemispheric ( S1-M1) connections in effective sensory-motor integration.

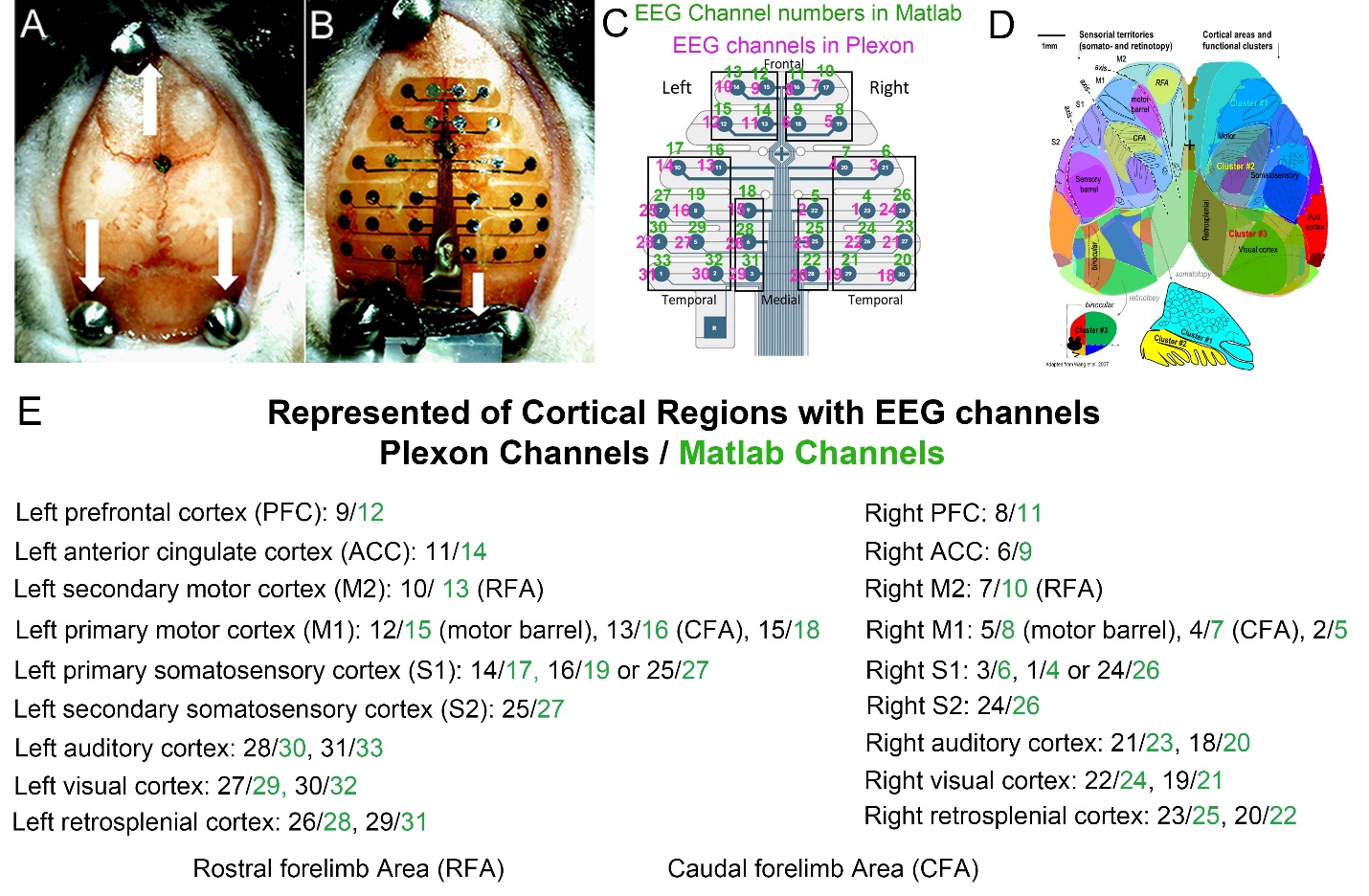

**Supplementary Figure 12: Overview of EEG Electrode Placement and Cortical Region Representation for MEA Recordings.** (A) Displays the skull post-surgery before planting the MEA probe (adapted from Jonak et al., 2018). Three screws are placed on the skull, with the reference electrode soldered to the screw on the left cerebellum. The Bregma is marked to align the probe accurately. (B) Details the electrode array layout on the skull (adapted from Jonak et al., 2018). The probe is positioned to align with the Bregma and skull sutures, ensuring consistent placement across subjects. (C) Shows the probe map with numbers in different systems (Plexon and Matlab), facilitating precise data analysis. (D) Adapted from Vanni etal., 2017, this panel maps the different cortical regions, serving as a reference for assigning cortical regions to the corresponding electrodes on the MEA probe. (E) Lists the specific brain areas represented by the electrode numbers in both Plexon and Matlab systems. This detailed mapping ensures accurate representation of the cortical regions monitored by each electrode, critical for robust EEG signal acquisition and analysis.
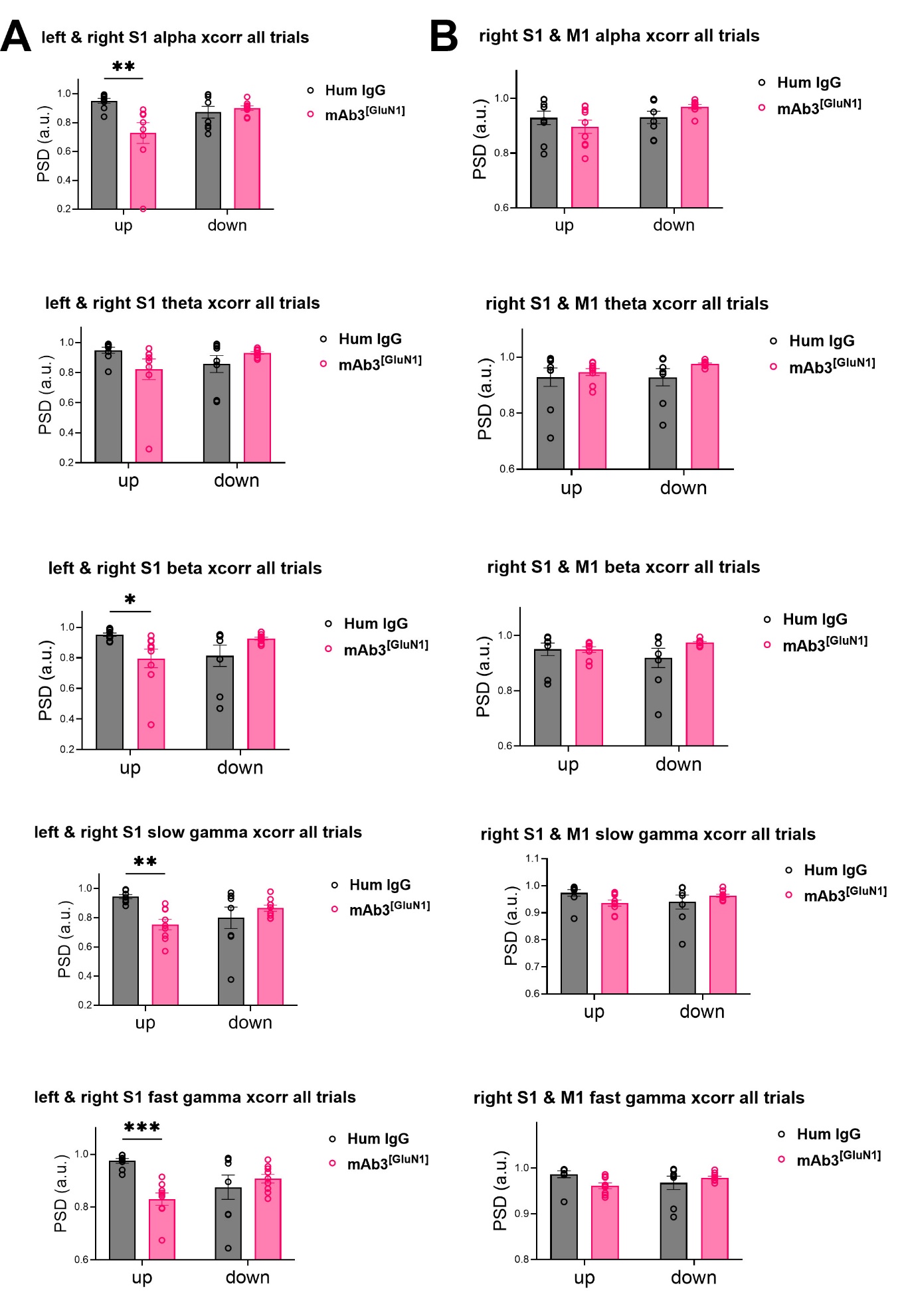

**Supplementary Figure 13: Analysis of inter-hemispheric (left-right S1) and intra-hemispheric (right S1-M1) functional connectivity in mAb3^[GluN1]^-treated male mice**. (A) Inter-hemispheric functional connectivity between left and right S1 across different frequency bands during all trials. 'up' corresponds to the EEG signals recorded when mice were performing the facing-up pole test, while 'down' refers to EEG signals during the facing-down pole test. Hum IgG was used as the control. In the 'up' trials, the control group showed significantly higher inter-hemispheric S1 connectivity than the mAb3^[GluN1]^ group across the alpha (P=0.0016), beta (P=0.037), slow gamma (P=0.0039), and fast gamma bands (P=0.0007). No significant difference was detected in the 'down' trials.

(B) Intra-hemispheric functional connectivity between right S1 and M1 (the antibody injection side) across different frequency bands during all trials. There was no significant difference in the S1-M1 functional connectivity between the control and mAb3^[GluN1]^ groups during either 'up' or 'down' trials across any frequency band. Control up trial (n=9); Control down trial (n=8); mAb3^[GluN1]^ up trial (n=9); mAb3^[GluN1]^ down trial (n=9).

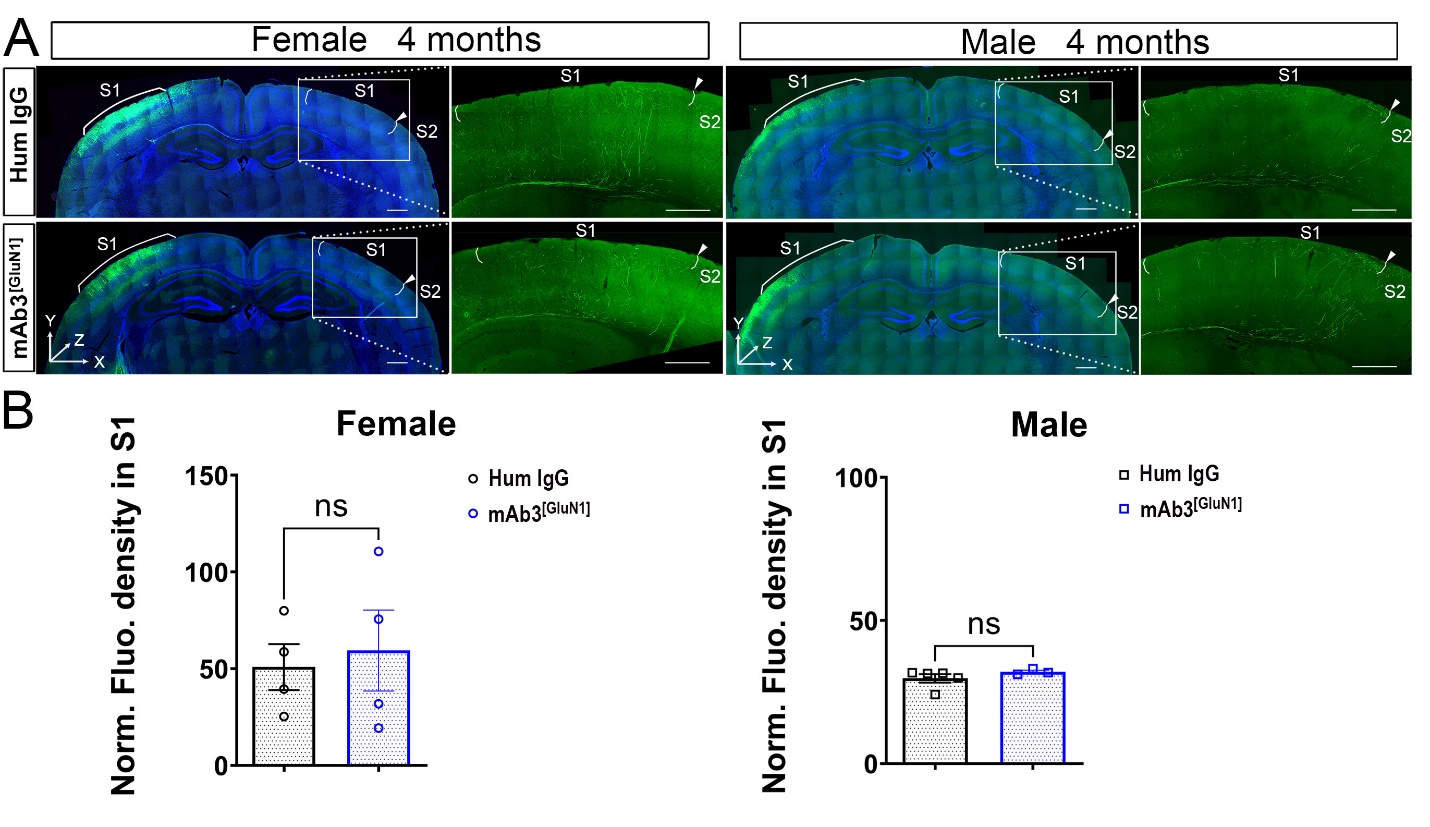

**Supplementary Figure 14: Recovery of S1 callosal projection density at 4 months following early mAb3^[GluN1]^ treatment.** (A) Mice treated with mAb3^[GluN1]^ from P3 to P12 show a recovery in S1 callosal projection density by 4 months of age, indicating reversion to typical levels for both genders. Human IgG treatment serves as the control. (B) Quantitative fluorescence density analysis reveals no significant difference in S1 callosal projection density between mAb3^[GluN1]^-treated and human IgG-treated controls at 4 months for both genders (P > 0.999 (female); P = 0.87 (male)), suggesting a normalization of early developmental alterations. n = 3 to 5 per group. Scale bar: 500μm.

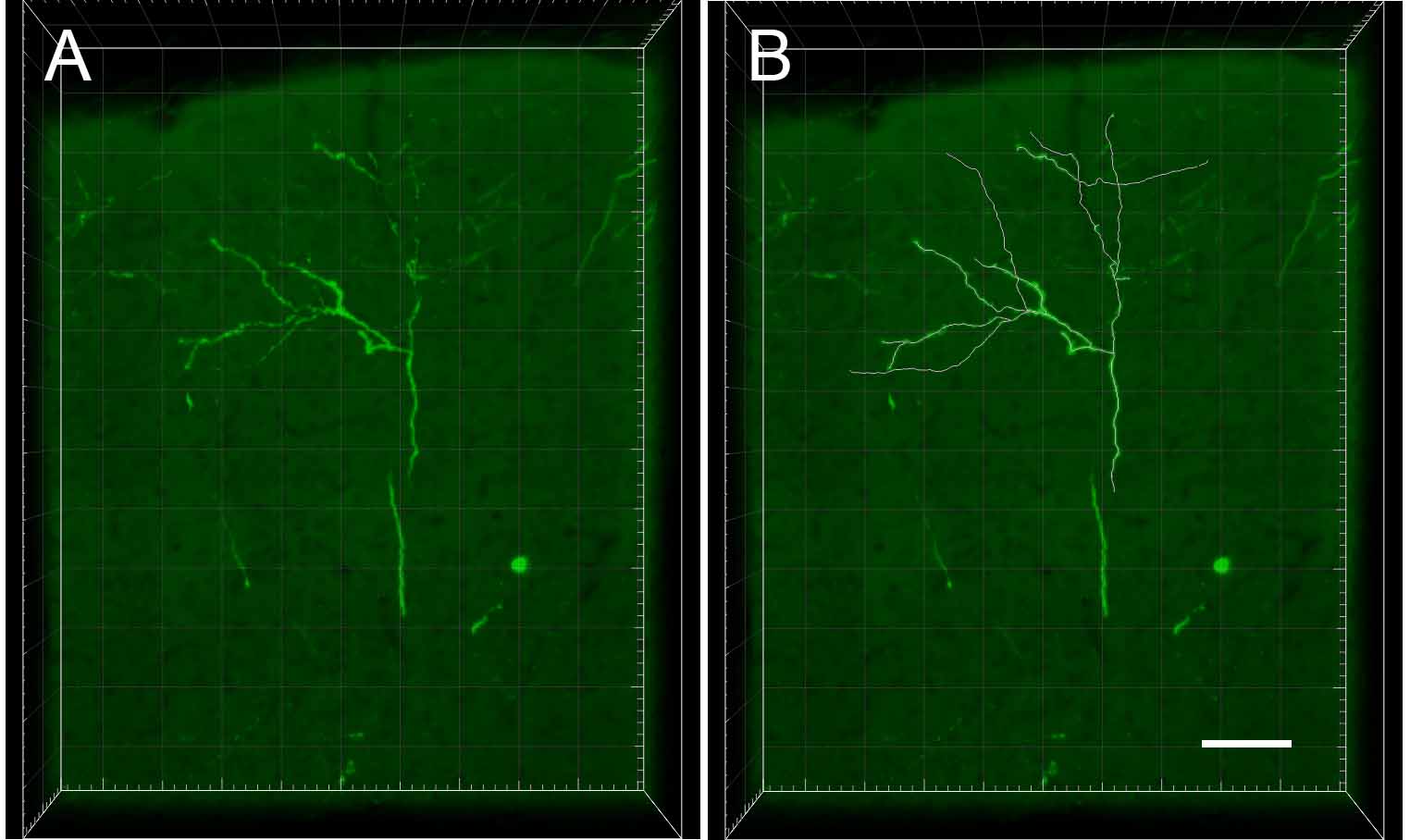

**Supplementary Figure 15: Axon branches were traced semi-automatically using Imaris software.** (A) 3D confocal image of S1 callosal axon terminals loaded in Imaris. Image was taken from a 200-μm coronal section of P120 mouse brain. (B) The morphology of individual callosal axon terminals was tracked across 3D in Imaris. Traced axon branches in Imaris. The morphology was tracked manually with combining Autopath function in Imaris. Scale bar: 70μm.

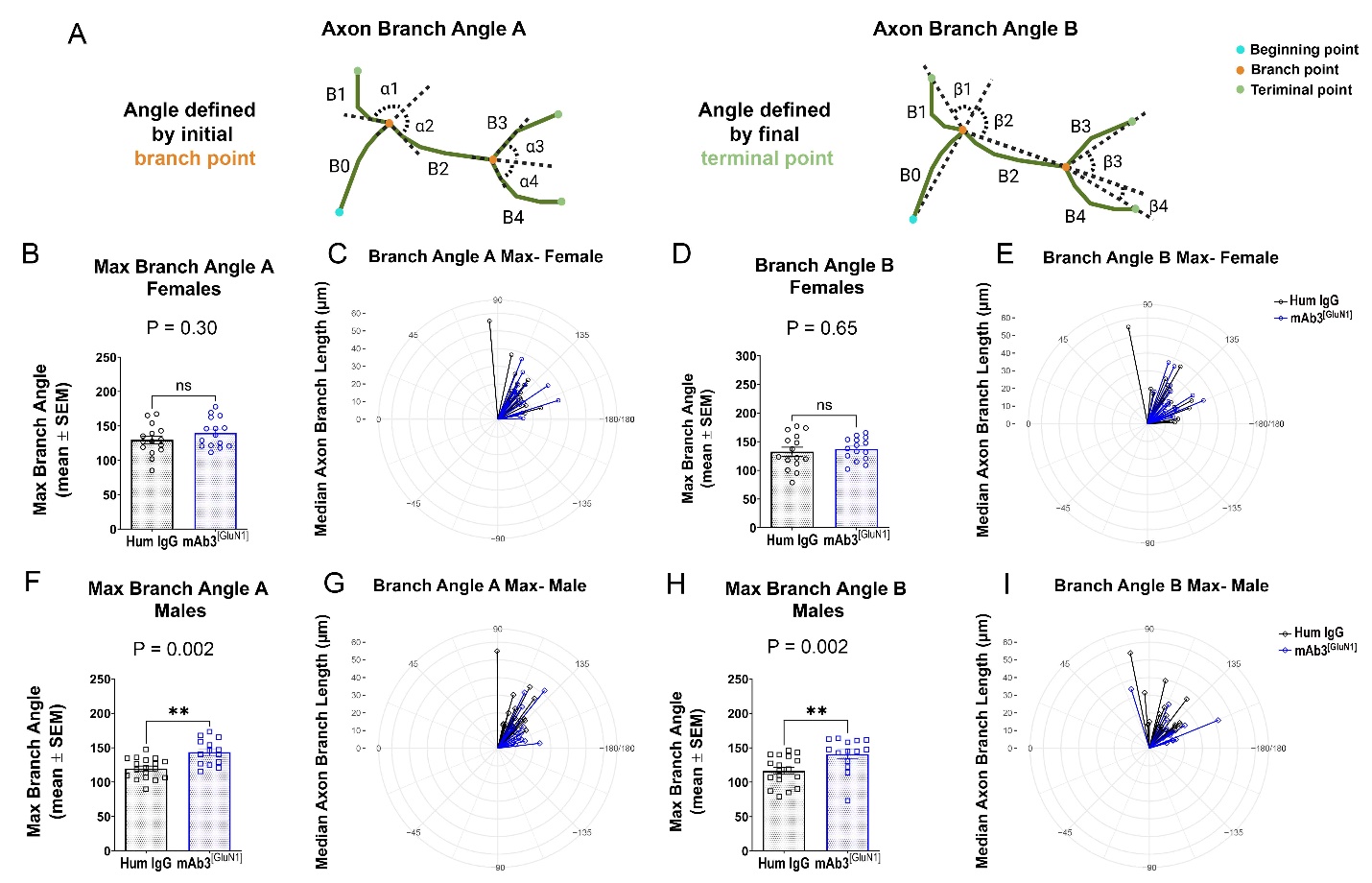

**Supplementary Figure 16: Significantly increased branch angles in mAb3^[GluN1]^-treated male mice.** (A) Diagram of two ways to measure axon branch angles. Axon branch angle A: defined as the angle between the extending lines from the branch point and its peripheral neighbor vertex. Axon branch angle B: defined as the angle between the extending lines connecting the branch point with the neighboring branch points and the terminal points, respectively. Max branch angle A in female (B, C) and male (F, G) mice. Max branch angle B in female (D, E) and male (H, I) mice. There were significantly increased branch angles in mAb3^[GluN1]^ male mice (F, H) in the two different axon branch angle measurements. n = 14 to 18 per group. The plots in C, E, G, and I were made in R using ggplot2 package. The p value on the graph represents the statistical difference between the two groups by using Mann-Whitney test.

**Supplementary** **Video 1: Represented balance beam performance of Human IgG treated female in 0.25x speed (Trial 4).**

**Supplementary** **Video 2: Represented balance beam performance of mAb3^[GluN1]^-treated female in 0.25x speed (Trial 4).**

**Supplementary** **Video 3: Represented balance beam performance of Human IgG treated male in 0.25x speed (Trial 4).**

**Supplementary** **Video 4: Represented balance beam performance of mAb3^[GluN1]^-treated male in 0.25x speed (Trial 4).**

**Supplementary** **Video 5: Represented facing up pole performance of Human IgG treated female in 0.5x speed (Trial 2).**

**Supplementary** **Video 6: Represented facing up pole performance of mAb3^[GluN1]^-treated female in 0.5x speed (Trial 2).**

**Supplementary** **Video 7: Represented facing up pole performance of Human IgG treated male in 0.5x speed (Trial 3).**

**Supplementary** **Video 8: Represented facing up pole performance of mAb3^[GluN1]^-treated male in 0.5x speed (Trial 3).**
